## Supplemental data for "Examining multiple cellular pathways at once using multiplex hextuple luciferase assaying"

**Supplementary Material**

**Supplementary Figures**

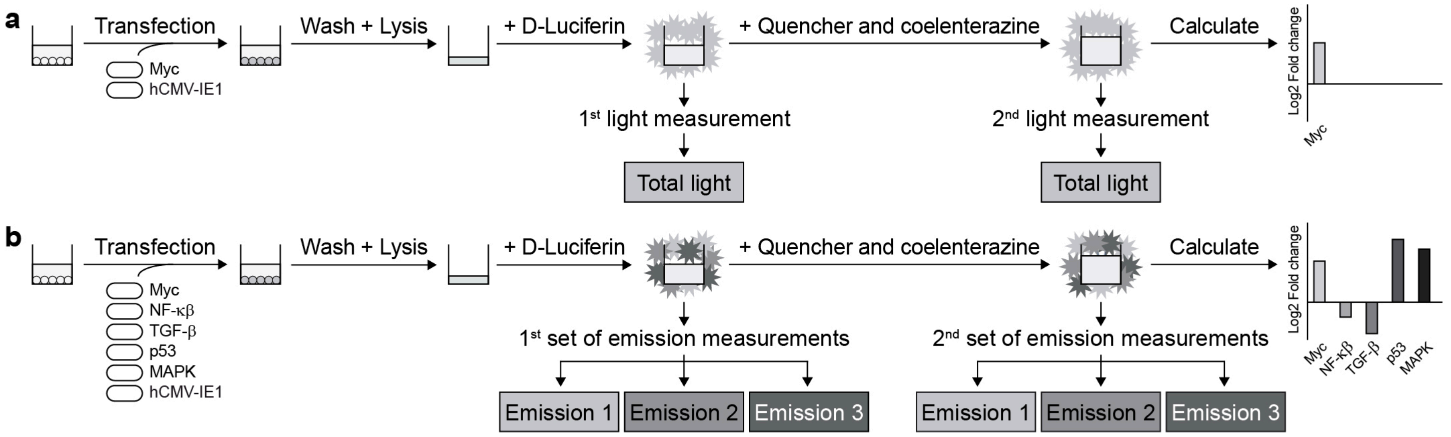


**Supplementary Figure 1. Development of multiplex luciferase assays for simultaneous analysis of cellular signaling events.** (**a**) Schematic of the dual-luciferase assay widely used for monitoring the activity of a single experimental cellular signaling event (e.g., transcriptional influence through c-Myc response elements) coupled to one luciferase, whose activity is normalized against a control cellular signaling event (e.g., constitutive hCMV-IE1 promoter) coupled to a second luciferase. A cell sample, previously co-transfected with experimental and control luciferase reporter plasmids, is washed and lysed before (**1**) addition of D-Luciferin substrate and measurement of its total light emission as an indicator of experimental cellular signaling, and (**2**) addition of a quenching reagent (to neutralize the first substrate-induced light emission) plus coelenterazine substrate, for which total light emission is measured as an indicator for control cellular signaling. Both light measurements are subsequently used to quantitate the experimental cellular signaling event (e.g., c-Myc signaling). (**b**) Simplified schematic of one possibility for a multiplex hextuple luciferase assay that can be used to simultaneously monitor five experimental cellular signaling events (e.g., transcriptional response signaling through c-Myc, NF-κβ, TGF-β, p53, and MAPK/JNK response elements). Each experimental parameter is coupled to a luciferase with a unique combination of substrate and spectral emission properties and normalized against a control cellular signaling event (e.g., hCMV-IE1 promoter). A cell sample, previously co-transfected with all experimental and control luciferase reporter plasmids, is washed and lysed before (**1**)

addition of D-Luciferin substrate and measurement of the first set of spectrally distinguishable emissions, and (**2**) addition of quenching reagent plus coelenterazine substrate and measurement of the resulting emissions. All six emission measurements are subsequently used to quantitate all five experimental cellular signaling events simultaneously (e.g., c-Myc, NF-κβ, TGF-β, p53, and MAPK/JNK signaling).

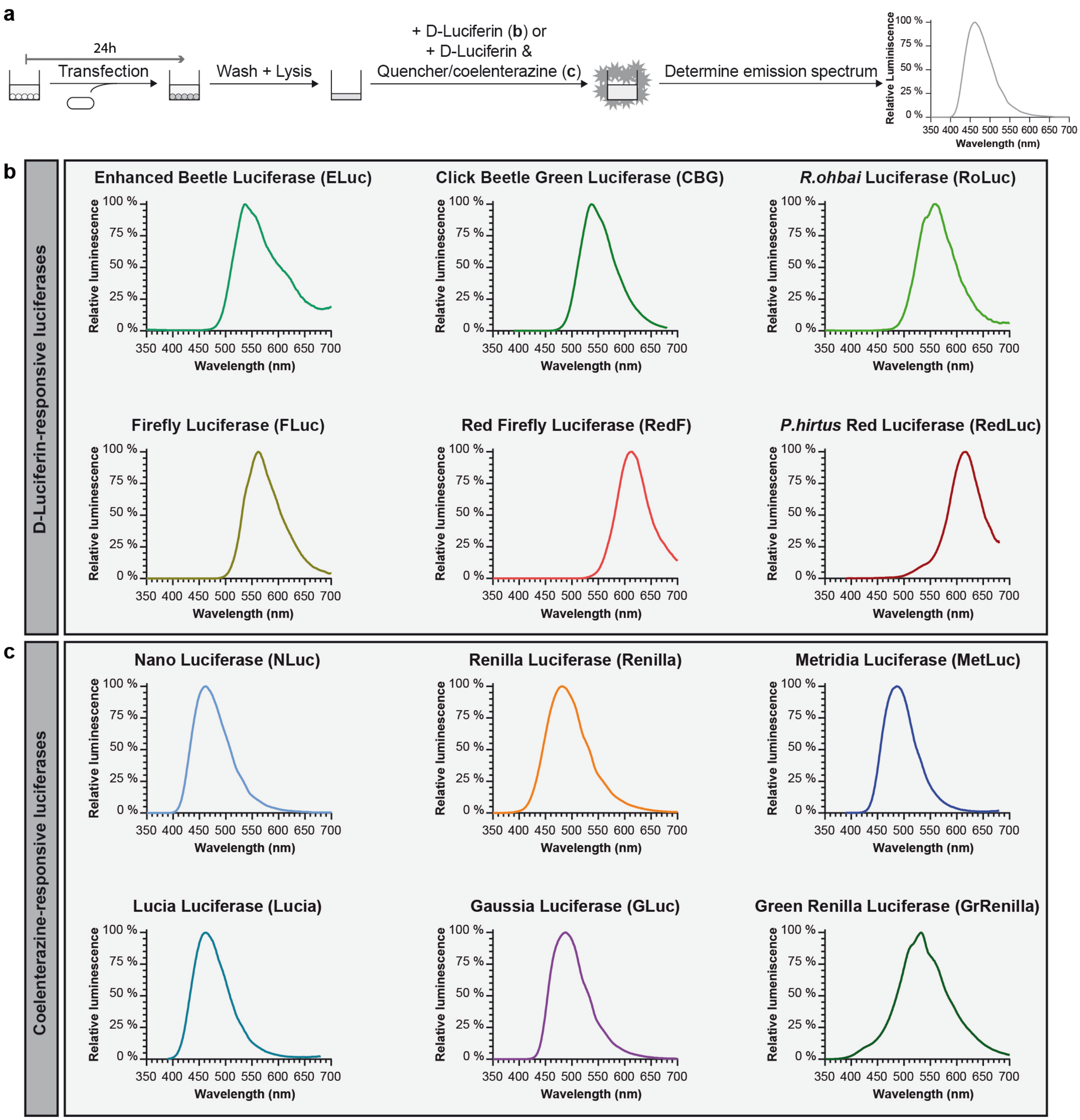


**Supplementary Figure 2. Emission spectra for the 12 luciferases examined in this study.** (**a**) Schematic of the experimental approach used to determine the emission spectrum for each luciferase: a plasmid encoding a single constitutively expressed luciferase transcriptional unit was transfected into HEK293T/17 cells. Cells were lysed 24 hours later and, after addition of the appropriate substrate (D-Luciferin alone or followed by addition of quencher/coelenterazine), the emission spectrum for each sample was recorded between 350 and 700 nm using the Linear Variable Filter emission monochromator of the CLARIOStar multimode microplate reader. (**b**) Emission spectra for the D-Luciferin-responsive luciferases. Because the Enhanced Beetle Luciferase (ELuc) exhibited a reduced emission intensity, a broader spectral bandwidth of 20 nm was applied to ensure enough light was captured during emission recording. (**c**) Emission spectra for the coelenterazine-responsive luciferases. The spectrum of NLuc was also recorded with its preferred substrate furimazine (data not shown); however, no apparent difference was observed when coelenterazine was used as the substrate.

**a**

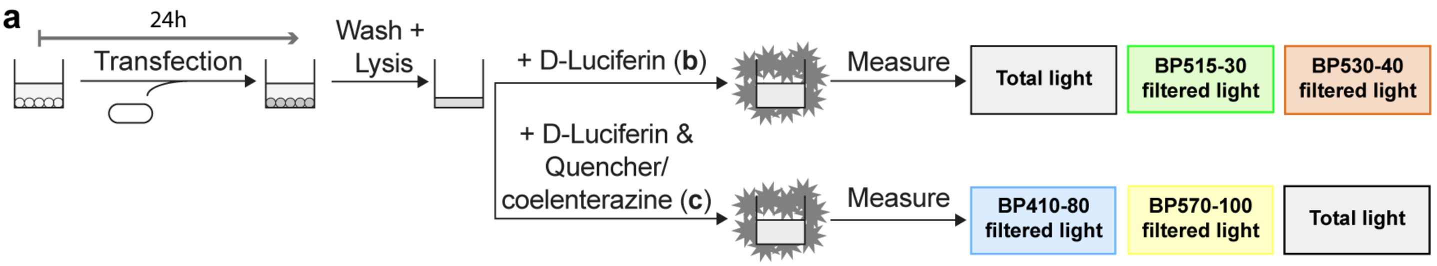


**b** **The calculation of transmission coefficients for D-Luciferin-responsive luciferases**

**BP515-30 filtered light**

$$\kappa Luciferase_{515} = \frac{Luciferase_{515}}{Luciferase_{TOTAL}}$$

**BP530-40 filtered light**

$$\kappa Luciferase_{530} = \frac{Luciferase_{530}}{Luciferase_{TOTAL}}$$

**c** **Transmission coefficients for D-Luciferin-responsive luciferases**

**BP515-30 filtered light**

$$\kappa ELuc_{515} = \frac{ELuc_{515}}{ELuc_{TOTAL}} = 24.32 \pm 0.08\%$$

$$\kappa FLuc_{515} = \frac{FLuc_{515}}{FLuc_{TOTAL}} = 7.25 \pm 0.01\%$$

$$\kappa RedF_{515} = \frac{RedF_{515}}{RedF_{TOTAL}} = 0.106 \pm 0.01\%$$

**BP530-40 filtered light**

$$\kappa ELuc_{530} = \frac{ELuc_{530}}{ELuc_{TOTAL}} = 46.15 \pm 0.26\%$$

$$\kappa FLuc_{530} = \frac{FLuc_{530}}{FLuc_{TOTAL}} = 29.80 \pm 0.04\%$$

$$\kappa RedF_{530} = \frac{RedF_{530}}{RedF_{TOTAL}} = 1.36 \pm 0.01\%$$

**d** **The calculation of transmission coefficients for coelenterazine-responsive luciferases**

**BP410-80 filtered light**

$$\kappa Luciferase_{410} = \frac{Luciferase_{410}}{Luciferase_{TOTAL}}$$

**BP570-100 filtered light**

$$\kappa Luciferase_{570} = \frac{Luciferase_{570}}{Luciferase_{TOTAL}}$$

**e** **Transmission coefficients for coelenterazine-responsive luciferases**

**BP410-80 filtered light**

$$\kappa NLuc_{410} = \frac{NLuc_{410}}{NLuc_{TOTAL}} = 14.62 \pm 0.12\%$$

$$\kappa Renilla_{410} = \frac{Renilla_{410}}{Renilla_{TOTAL}} = 4.46 \pm 0.07\%$$

$$\kappa GrRenilla_{410} = \frac{GrRenilla_{410}}{GrRenilla_{TOTAL}} = 1.94 \pm 0.27\%$$

**BP570-100 filtered light**

$$\kappa NLuc_{570} = \frac{NLuc_{570}}{NLuc_{TOTAL}} = 12.87 \pm 0.13\%$$

$$\kappa Renilla_{570} = \frac{Renilla_{570}}{Renilla_{TOTAL}} = 27.93 \pm 0.07\%$$

$$\kappa GrRenilla_{570} = \frac{GrRenilla_{570}}{GrRenilla_{TOTAL}} = 57.42 \pm 0.38\%$$

**Supplementary Figure 3. Determination of transmission coefficients for each luciferase over the empirically-determined bandpass emission filters.** (**a**) Schematic of the experimental setup performed to determine the transmission coefficients for each luciferase using absolute luminescence *in toto* or over the indicated bandpass emission filters. (**b**) The transmission coefficients (**κ**) of each D-Luciferin-responsive luciferase over the indicated bandpass emission filters. **κ**$Luciferase_{515}$ and **κ**$Luciferase_{530}$ were calculated by dividing the light that was transmitted for each luciferase through each of the filters, $Luciferase_{515}$ and $Luciferase_{530}$, respectively, by the total light emitted by each luciferase ($Luciferase_{TOTAL}$). (**c**) For the three D-Luciferin-responsive luciferases, **κ**$ELuc_{515}$, **κ**$FLuc_{515}$, and **κ**$RedF_{515}$ represent the transmission coefficients over the BP515-30 bandpass emission filter (**Left**), while **κ**$ELuc_{530}$, **κ**$FLuc_{530}$, and **κ**$RedF_{530}$ represent the transmission coefficients over the BP530-40 bandpass emission filter (**Right**). (**d**) The transmission coefficients (**κ**) of each coelenterazine-responsive luciferase over the indicated bandpass emission filters, **κ**$Luciferase_{410}$ and **κ**$Luciferase_{570}$, were calculated by dividing the light that was transmitted for each luciferase through each of the filters, $Luciferase_{410}$ and $Luciferase_{570}$, respectively, by the total light emitted by each luciferase ($Luciferase_{TOTAL}$). (**e**) For the three coelenterazine-responsive luciferases, **κ**$NLuc_{410}$, **κ**$Renilla_{410}$, and **κ**$GrRenilla_{410}$ represent the transmission coefficients over the BP410-80 bandpass emission filter (**Left**), while **κ**$NLuc_{570}$, **κ**$Renilla_{570}$, and **κ**$GrRenilla_{570}$ represent the transmission coefficients over the BP570-100 bandpass emission filter (**Right**). Mathematical tools to perform and facilitate these calculations are provided as a protected Microsoft Excel file (**Supplementary Figure 21**).

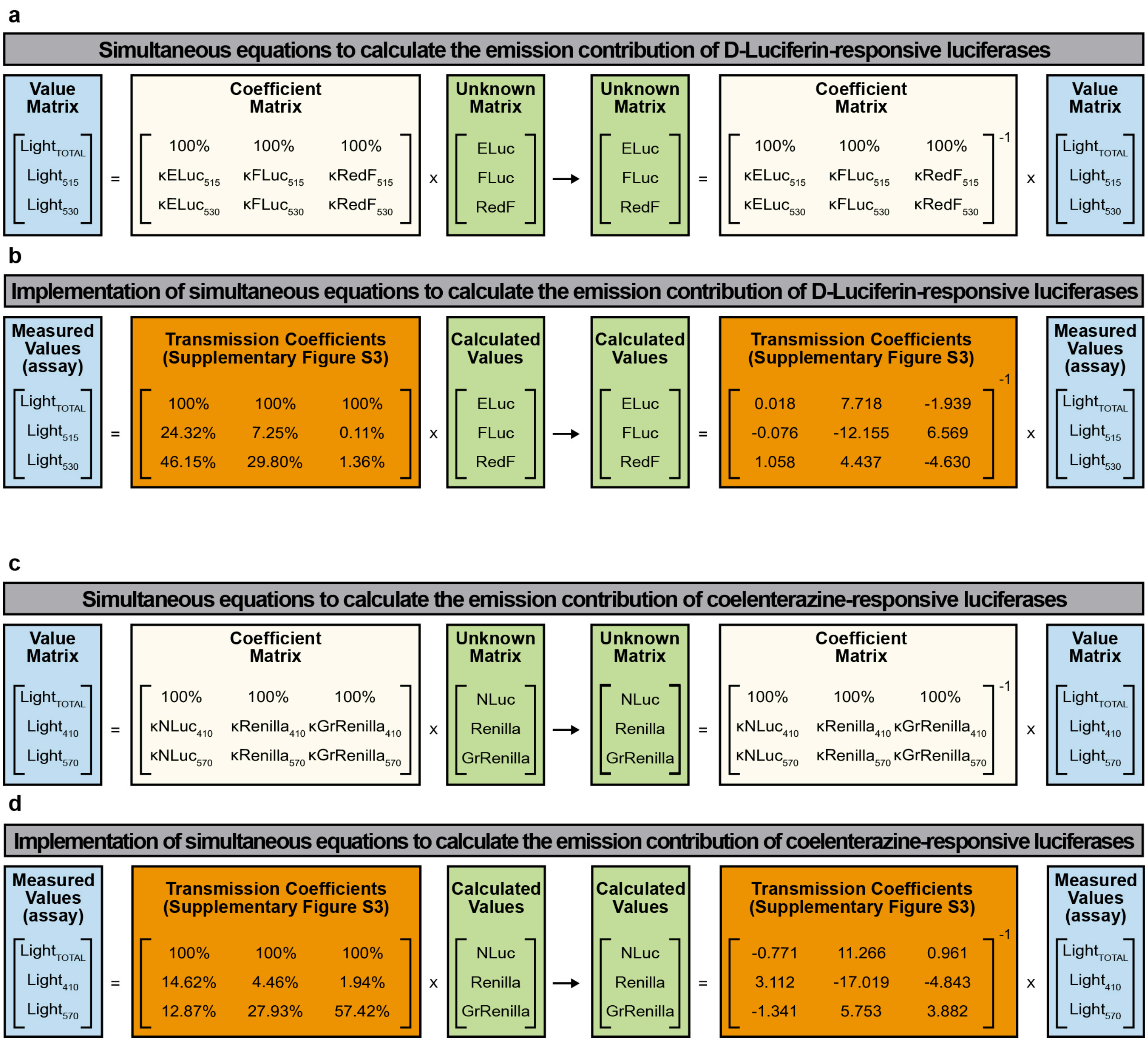


**Supplementary Figure 4. Calculation of emission contributions of individual luciferases in a mixture of three luciferases using simultaneous equations.** (**a**) Simultaneous equations solving for the D-Luciferin-responsive luciferase contributions have three unknowns corresponding to the amount of each D-Luciferin-responsive luciferase in a mix, namely ELuc, FLuc, and RedF. The value matrix includes the three measured values for the first step of the luciferase assay, while the coefficient matrix includes all the transmission coefficients for the luciferases in the equation system. To solve the simultaneous equations for the D-Luciferin-responsive luciferases (unknown matrix), the inverse of the coefficient matrix was multiplied by the value matrix. $Light_{TOTAL}$, $Light_{515}$, and $Light_{530}$ represent the total measured light values and the light filtered by the BP515-30 and BP530-40 bandpass emission filters for the D-Luciferin-responsive luciferases (ELuc, FLuc, and RedF). (**b**) To obtain calculated values for each D-Luciferin-responsive luciferase-linked reporter unit, a matrix inversion of the coefficient matrix (the matrix containing values for all transmission coefficients) previously obtained using the appropriate bandpass emission filters (**Supplementary Figure 3b** and **c**), was multiplied by the value matrix (the matrix containing luminescence measurements obtained by the plate reader). (**c**) Simultaneous equations solving for the coelenterazine-responsive luciferase contributions have three unknowns corresponding to the amount of the coelenterazine-responsive luciferase in a mix, namely NLuc, Renilla, and GrRenilla. The value matrix includes the three measured values for the second step of the luciferase assay, while the coefficient matrix includes all the transmission coefficients for the luciferases in the equation system. To solve the simultaneous equations for the coelenterazine-responsive luciferases (the unknown matrix), the inverse of the coefficient matrix was multiplied by the value matrix. $Light_{TOTAL}$, $Light_{410}$, and $Light_{570}$ represent the total measured light values and the light filtered by the BP410-80 and BP570-100 bandpass emission filters for the coelenterazine-responsive luciferases (NLuc, Renilla, and GrRenilla). (**d**) To obtain calculated values for each coelenterazine-responsive luciferase-linked reporter unit, a matrix inversion of the coefficient matrix (the matrix containing values for all transmission coefficients) previously obtained using the appropriate bandpass emission filters (**Supplementary Figure 3d** and **e**), was multiplied by the value matrix (the matrix containing luminescence measurements obtained by the plate reader). Mathematical tools and guidelines to perform and facilitate these calculations are provided as protected Microsoft Excel file (**Supplementary Figure 22, Supplementary Figure 23**, and **Supplementary Figure 24**).

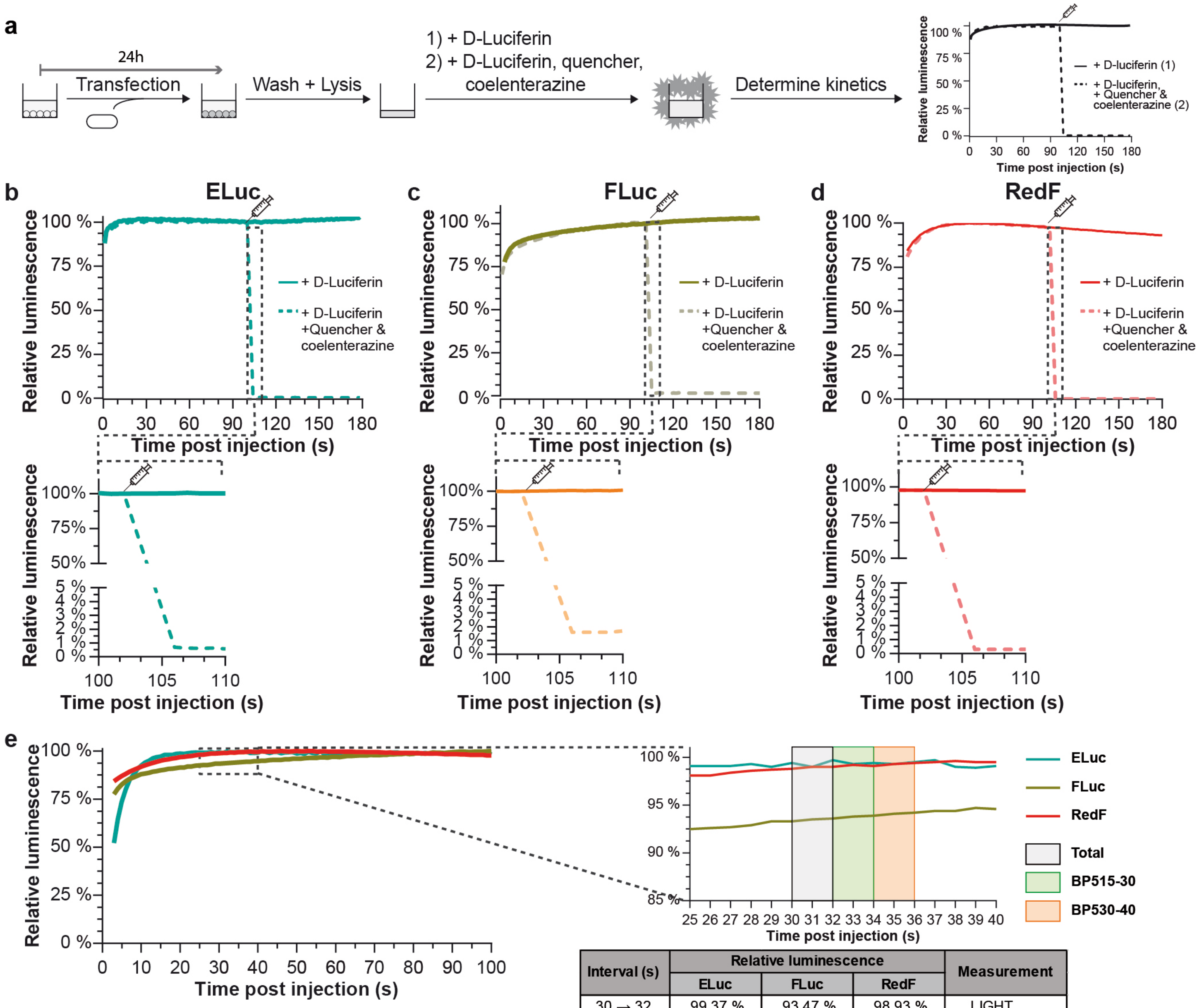


| Interval (s) | Relative luminescence | | | Measurement |
|---|---|---|---|---|
| | ELuc | FLuc | RedF | |
| 30 → 32 | 99.37 % | 93.47 % | 98.93 % | $LIGHT_{TOTAL}$ |
| 32 → 34 | 99.47 % | 93.77 % | 99.10 % | $LIGHT_{515}$ |
| 34 → 36 | 99.40 % | 94.07 % | 99.27 % | $LIGHT_{530}$ |

**Supplementary Figure 5. Determination of kinetic parameters for ELuc, FLuc, and RedF luciferases during the first step of the multiplex hextuple luciferase assay.** (**a**) Experimental setup to determine the glowing and quenching kinetics of the three D-Luciferin-responsive luciferases, ELuc, FLuc, and RedF. After transfection and a 24-hour incubation, cells were washed and lysed before D-Luciferin substrate buffer (LARII buffer) was added. Then the reaction was monitored for 180 seconds with 1 second measurements taken every second to determine the emission kinetics (**thick line**). In a duplicate parallel reaction, quencher and coelenterazine substrate buffer (Stop & Glo buffer) were added at 100 seconds and the decaying luminescence was measured to determine the quenching kinetics (**dashed line**). (**b-d**) Glowing and quenching kinetics of the three D-Luciferin luciferases, ELuc (**b**), FLuc (**c**), and RedF (**d**) over a 180-second interval (**Top**) and a 10-second interval (**Bottom**) to illustrate acute quenching kinetics after the addition of quencher and coelenterazine. (**e**) Determination of the time interval to perform emission measurements during the first step (after the addition of the D-Luciferin-containing LARII buffer) of the multiplex luciferase assay. Overlay of the kinetic charts of ELuc, FLuc, and RedF (**Left**), and a close-up view of the section between 25 and 40 seconds (**Right**). Two bandpass emission filters, one between 500 and 530 nm (BP515-30) and another between 510 and 550 nm (BP530-40), were used to capture the maximum amount of light emitted by ELuc and FLuc (**Figure 1d** and **Figure 1g**), respectively. Luminescence is represented in relative units. The amount of light recorded as relative luminescence, which was captured during each of the three most stable 2-second intervals ($Light_{515}$, $Light_{530}$, and $Light_{TOTAL}$), is shown below the graph. Source data are provided as a Source Data file.

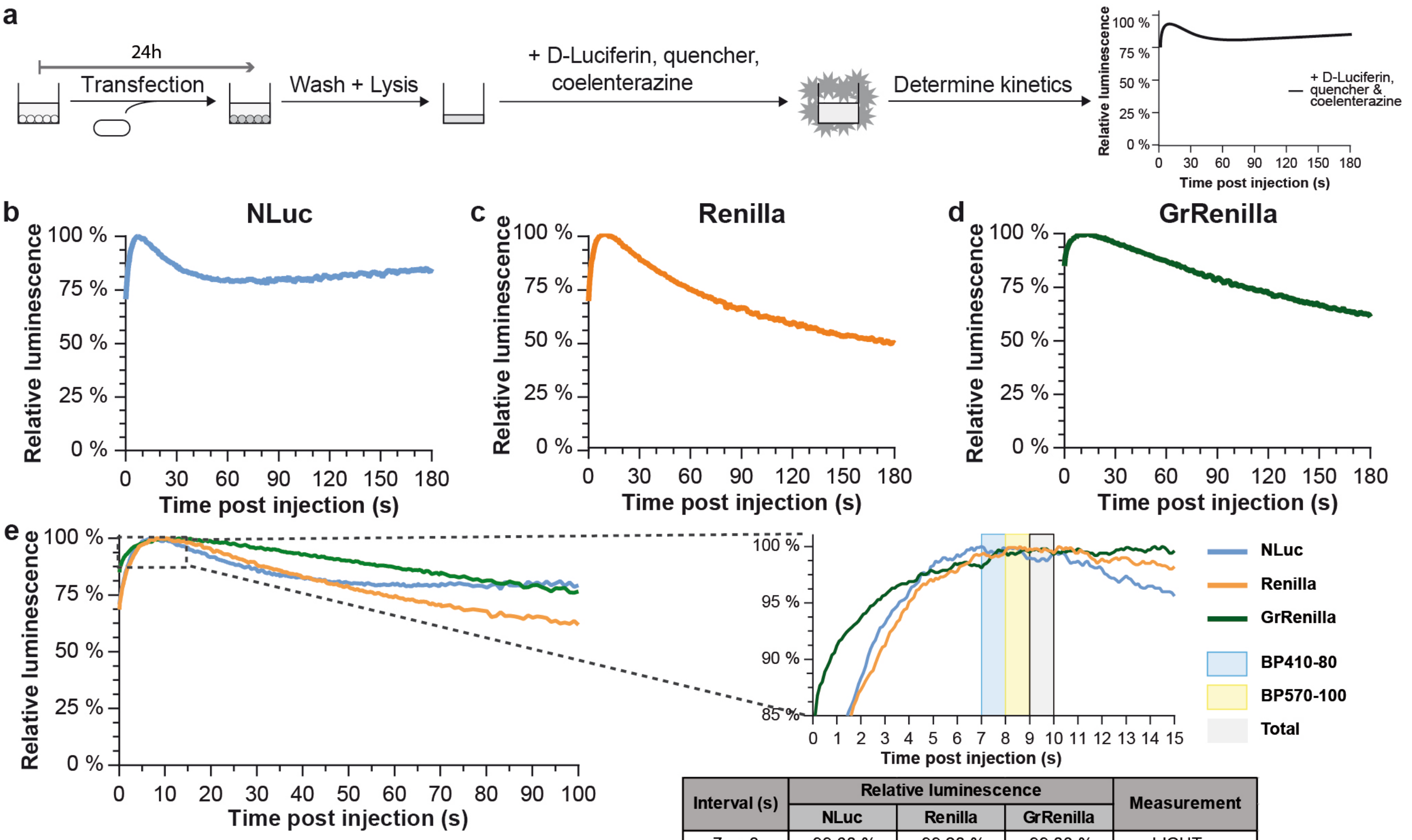


| Interval (s) | Relative luminescence | | | Measurement |
|---|---|---|---|---|
| | NLuc | Renilla | GrRenilla | |
| 7 → 8 | 99.60 % | 99.20 % | 99.03 % | $LIGHT_{410}$ |
| 8 → 9 | 99.62 % | 99.83 % | 99.47 % | $LIGHT_{570}$ |
| 9 → 10 | 98.88 % | 99.70 % | 99.68 % | $LIGHT_{TOTAL}$ |

**Supplementary Figure 6. Determination of the kinetic parameters for NLuc, Renilla, GrRenilla during the second step of the multiplex hextuple luciferase assay.** (**a**) Experimental setup used to determine the flash kinetics of the three coelenterazine-responsive luciferases, NLuc, Renilla, and GrRenilla. After transfection and a 24-hour incubation, cells were washed and lysed before D-Luciferin substrate buffer (LARII buffer), as well as a quencher and coelenterazine substrate buffer (Stop & Glo buffer), were added. Then the reaction was monitored over 180 seconds with 0.1-second measurements taken every 0.1 seconds to determine the fast-changing flash kinetics. (**b-d**) Determination of flash kinetics of the three coelenterazine-responsive luciferases, NLuc (**b**), Renilla (**c**), and GrRenilla (**d**). (**e**) Determination of the time interval to perform emission measurements during the second step of the multiplex luciferase assay (after the addition of luciferase-quenching agent and coelenterazine substrate-containing Stop & Glo buffer). Overlay of the kinetics of NLuc, Renilla, and GrRenilla and a close-up view of the section between 0 and 15 seconds. Two bandpass emission filters, one between 370 and 450 nm (BP410-80) and another between 520 and 620 nm (BP570-100), were used to capture the maximum light emitted by NLuc and GrRenilla, respectively (**Figure 1f** and

**Figure 1h**). Luminescence is represented in relative units. The amount of light recorded as relative luminescence captured during each of the three most stable 1-second intervals ($Light_{410}$, $Light_{570}$, and $Light_{TOTAL}$), is shown below the graph. Source data are provided as a Source Data file.

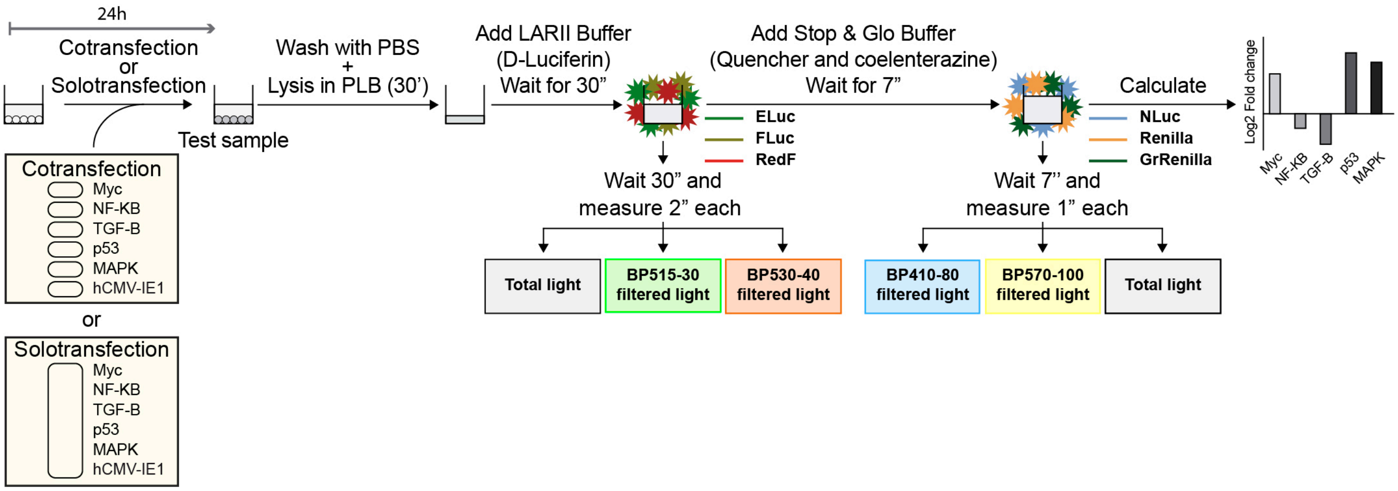


**Supplementary Figure 7. Detailed overview of the empirically-determined multiplex luciferase assay.** A cell sample is washed with phosphate buffered saline (PBS) at 24 hours after cotransfection or solotransfection, followed by lysis for 30 minutes using the Promega Passive Lysis Buffer (PLB). The sample is then transferred to a plate reader equipped with the appropriate bandpass emission filters. Next, D-Luciferin-containing substrate buffer (Promega LARII buffer) is added and then three emission measurements are recorded for two seconds each starting at 30 seconds later: total light, BP515-30-filtered light, and BP530-40-filtered light (**Figure 1g**). Finally, a D-Luciferin luciferase quencher and coelenterazine-containing substrate buffer (the Promega Stop & Glo buffer) are added and then three additional emission measurements are recorded at one second each starting 7 seconds later: total light, BP410-80-filtered light, and BP570-100-filtered light (**Figure 1h**).

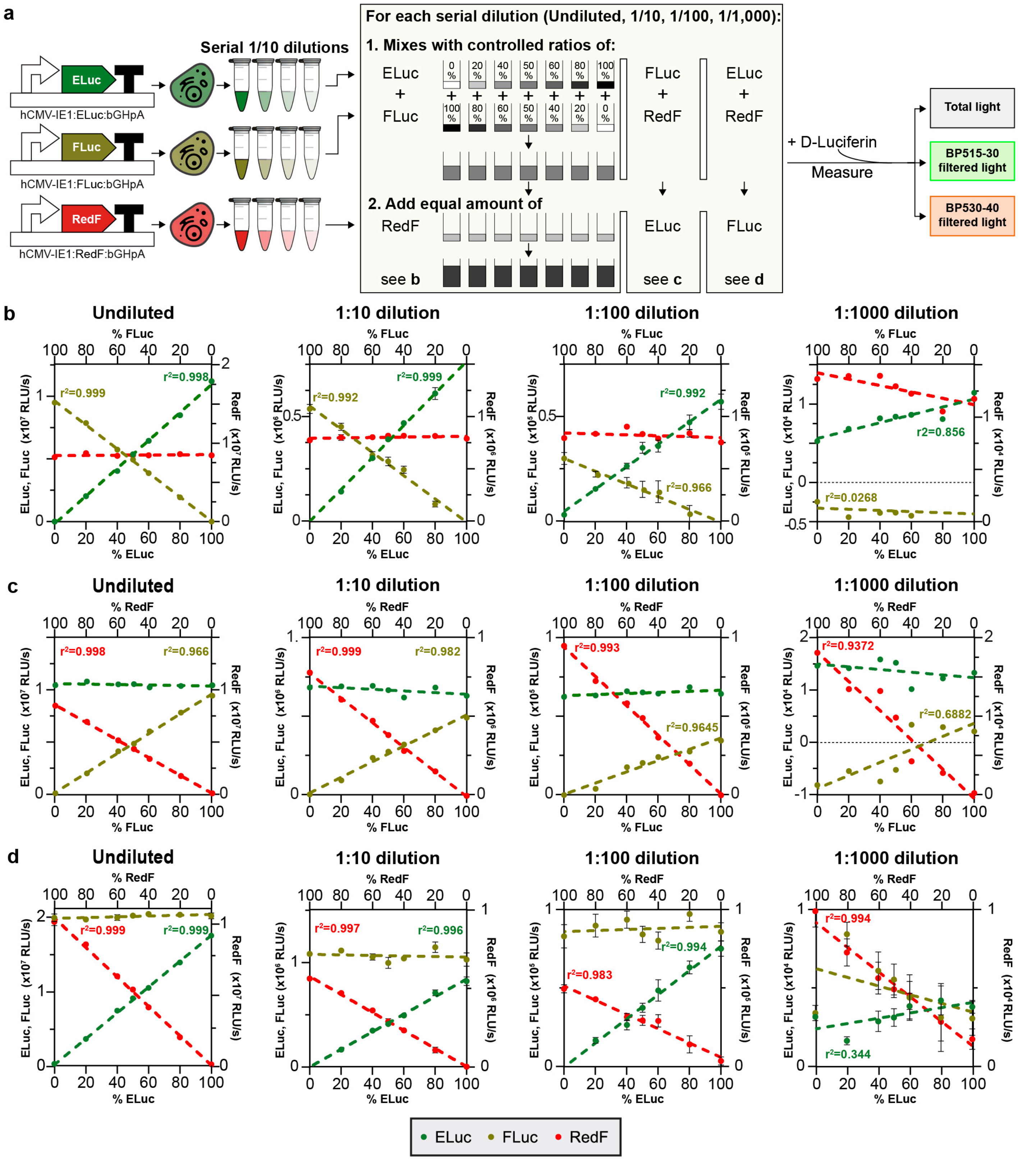


**Supplementary Figure 8. Determination of the dynamic range of the quantitative relationships between ELuc, FLuc, and RedF**. (**a**) Schematic of the experimental setup used to determine the dynamic range of the quantitative relationships between the D-Luciferin-responsive luciferases (ELuc, FLuc, and RedF) in a single emission recording experiment. Individual plasmids, each possessing one D-Luciferin-responsive transcriptional luciferase unit containing the hCMV-IE1 promoter, D-Luciferin-responsive luciferase, and transcriptional terminator were transfected into HEK293T/17 cells. After 24 hours, transfected cells were harvested, lysed, and 1:10 serial dilutions were prepared. Next, to determine the dynamic range of the quantitative relationship between ELuc and FLuc at a specific dilution of the serial dilution series, defined amounts of each were mixed at different ratios totaling 100%, followed by the addition of an equal amount of FLuc. After the addition of D-Luciferin substrate-containing buffer (LARII buffer), the total and filtered light were measured after 30 seconds (**Figure 1g**). Similar experimental setups were used to determine the dynamic range of the quantitative FLuc/RedF and ELuc/RedF relationships. (**b**) Determination of the dynamic range of the quantitative relationships between ELuc and FLuc at the indicated dilutions. The separation of the D-Luciferin-responsive luciferases was successful in a dynamic range from $10^7$ to $10^5$ RLU/s (down to a 1:100 dilution of the original lysate), but not $10^4$ RLU/s (1:1,000 dilution). The maximum rate measured was $10^7$ RLU/s. Similar results were observed with FLuc and RedF (**c**) and ELuc and RedF (**d**). For **b-d**, $P < 0.0001$ for all regression lines at varying concentrations, except for the largest dilutions (1:1,000 dilution) (right panels for **b-d**). For luciferases kept at constant concentrations, all minimal slopes interpolated by regression did not significantly differ from zero. Four technical replicates are included in each data point, and the standard error of the mean is represented. n=5 for c, 1:1000 dilution. Source data are provided as a Source Data file.

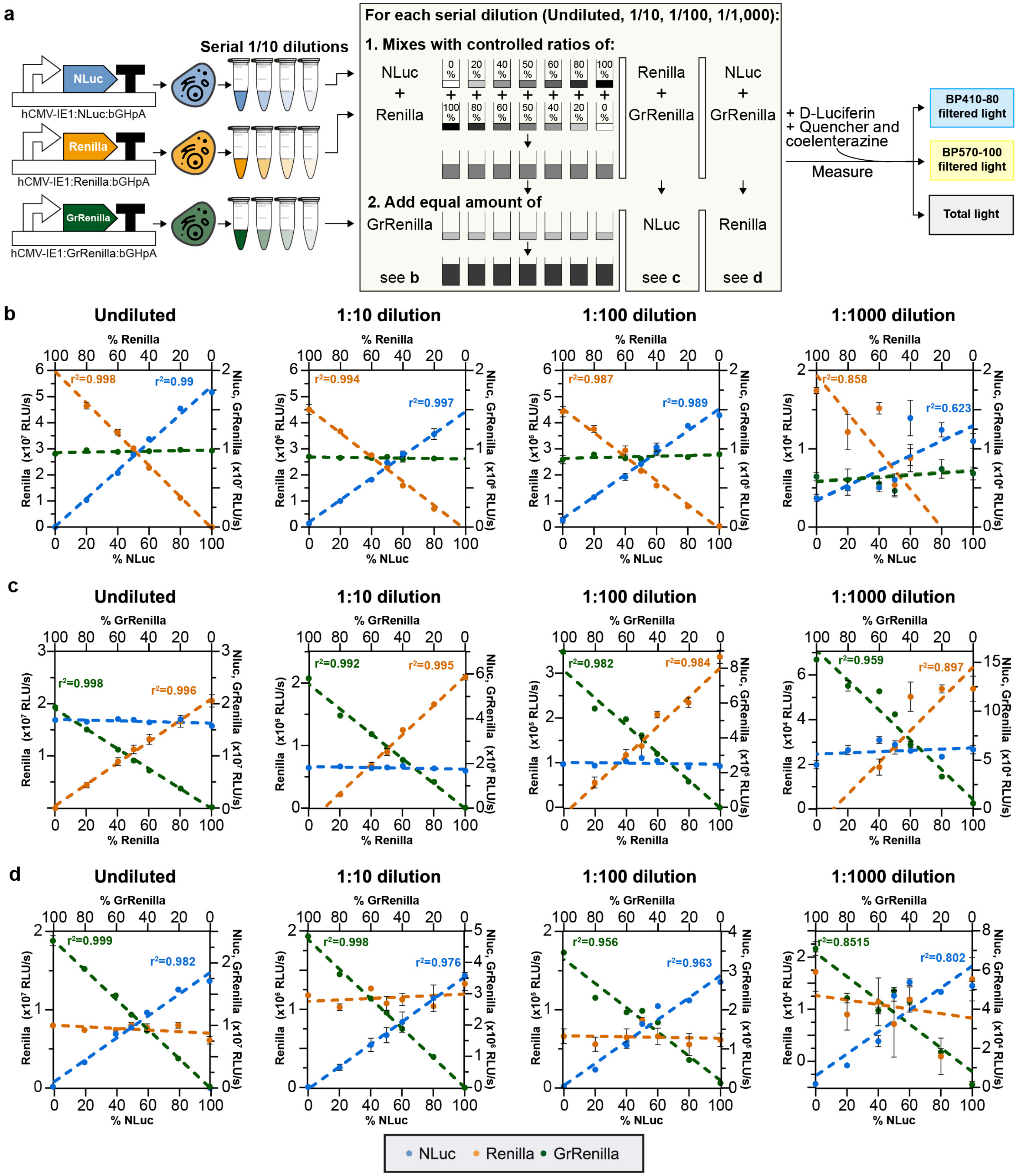


**Supplementary Figure 9. Determination of the dynamic range of the quantitative relationships between NLuc, Renilla, and GrRenilla.** (**a**) Schematic of the experimental setup used to determine the dynamic range of the quantitative relationships between the coelenterazine-responsive luciferases (NLuc, Renilla, and GrRenilla) in a single emission recording experiment. Individual plasmids, each possessing one transcriptional coelenterazine-responsive luciferase unit containing the hCMV-IE1 promoter, coelenterazine-responsive luciferase, and transcriptional terminator were transfected into HEK293T/17 cells. After 24 hours, transfected cells were harvested, lysed, and 1:10 serial dilutions were prepared. To determine the dynamic range of the quantitative relationship between NLuc and Renilla at a specific dilution of the serial dilutions series, defined amounts of each were mixed at different ratios totaling 100%, followed by the addition of an equal amount of GrRenilla. After the addition of D-Luciferin substrate-containing buffer (LARII buffer), as well as quencher and coelenterazine substrate-containing buffer (Stop & Glo buffer), total and filtered light were measured after 7 seconds (**Figure 1h**). Similar experimental setups were used to determine the dynamic range of the quantitative Renilla/GrRenilla and NLuc/GrRenilla relationships. (**b-d**) Determination of the dynamic range of the quantitative relationships between NLuc and Renilla for the dilutions shown. Separation of the coelenterazine-responsive luciferases was successful in a dynamic range from $10^7$ to $10^5$ RLU/s (down to a 1:100 dilution of the original undiluted lysate), but not $10^4$ RLU/s (1:1,000 dilution). The maximum rate measured was $10^7$ RLU/s. Similar results were observed when determining the dynamic range of the quantitative relationship between Renilla and GrRenilla (**c**) and NLuc and GrRenilla (**d**). For **b-d**, P <0.0001 for all regression lines at varying concentrations except for the largest dilutions (1:1,000 dilution) (right panel for **b-d**). For luciferases kept at constant concentrations, all minimal slopes interpolated by regression did not significantly differ from zero. Four technical replicates are included in each data point, and five for the 1:10000 dilution, and the standard error of the mean is represented. n=3 for d 1:10 dilution. Source data are provided as a Source Data file.

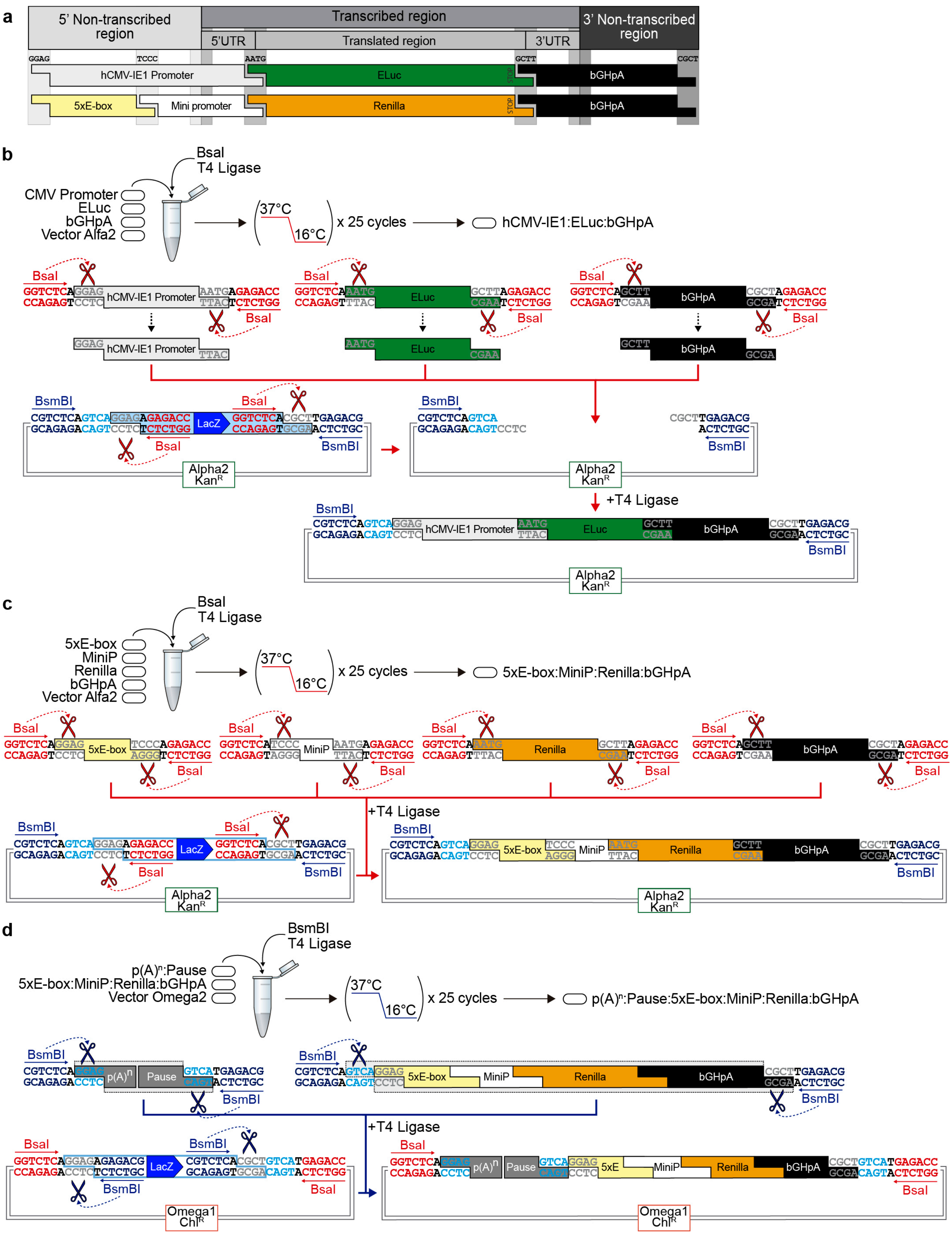
a
5' Non-transcribed region
Transcribed region
3' Non-transcribed region
5'UTR
Translated region
3'UTR
GGAG
TCCC
AATG
GCTT
CGCT
hCMV-IE1 Promoter
ELuc
bGHpA
5xE-box
Mini promoter
Renilla
bGHpA
b
BsaI
T4 Ligase
CMV Promoter
ELuc
bGHpA
Vector Alfa2
37°C
16°C
x 25 cycles
hCMV-IE1:ELuc:bGHpA
BsmBI
LacZ
Alpha2
KanR
+T4 Ligase
c
BsaI
T4 Ligase
5xE-box
MiniP
Renilla
bGHpA
Vector Alfa2
37°C
16°C
x 25 cycles
5xE-box:MiniP:Renilla:bGHpA
+T4 Ligase
d
BsmBI
T4 Ligase
p(A)n:Pause
5xE-box:MiniP:Renilla:bGHpA
Vector Omega2
37°C
16°C
x 25 cycles
p(A)n:Pause:5xE-box:MiniP:Renilla:bGHpA
+T4 Ligase
Omega1
ChlR

**Supplementary Figure 10. Schematic of the approach used to construct synthetic multipartite and binary assemblies to generate multi-luciferase reporter plasmids.** (**a**) Schematic illustrating the different synthetic assembly cloning overhangs (GoldenBraid 2.0 grammar) used to stitch the depicted DNA elements together to build the two transcriptional units illustrated in (**b**) and (**c**). The orthogonal 4-bp sequences GGAG, TCCC, AATG, GCTT, and CGCT are Type IIs restriction enzyme overhangs (BsaI) that allow directional assembly of pre-made DNA fragments into defined transcriptional units, as established by GoldenBraid2.0 rules (**Figure 3a** and **Figure 3b**). (**b**) Multipartite assembly of three pre-made DNA fragments (constitutive hCMV-IE1 promoter, coding sequence of ELuc luciferase, and bGH terminator) into the Alpha2 destination vector in a one-step, one-pot GoldenBraid 2.0 assembly reaction with BsaI and T4 ligase. Briefly, pre-made standard DNA fragments are digested by the Type IIs restriction enzyme BsaI and then the appropriate overhangs are ligated together into the destination vector, as established by GoldenBraid 2.0 rules, resulting in the constitutively expressed luciferase transcriptional unit (denoted as hCMV-IE1:ELuc:bGHpA) (**Figure 3c**). (**c**) Multipartite assembly of four pre-made DNA fragments (five copies of the E-box operator element, minimal promoter called MiniP, coding sequence of Renilla luciferase, and bGH terminator) into the Alpha2 destination vector in a one-step, one-pot GoldenBraid 2.0 assembly reaction with BsaI and T4 ligase. This assembly results in a pathway-responsive luciferase transcriptional unit (denoted as 5xEbox:Renilla:bGHpA) (**Figure 3d**). (**d**) Binary assembly of two components (a transcription blocker called $p(A)^n$:Pause and the pathway-responsive unit 5xEbox:Renilla:bGHpA) into the Omega2 destination vector in a one-step, one-pot GoldenBraid 2.0 assembly reaction with BsmBI and T4 ligase. This assembly results in an insulated c-Myc pathway-responsive luciferase unit. Assembled components in the complementary vectors (Alpha1 and Alpha2) are digested by the Type IIs restriction enzyme BsmBI and then ligated together as established by GoldenBraid 2.0 rules (**Figure 3e**). $Kan^R$ and $Chl^R$ represent kanamycin and chloramphenicol resistance markers, respectively, for bacterial selection of the DNA clones.

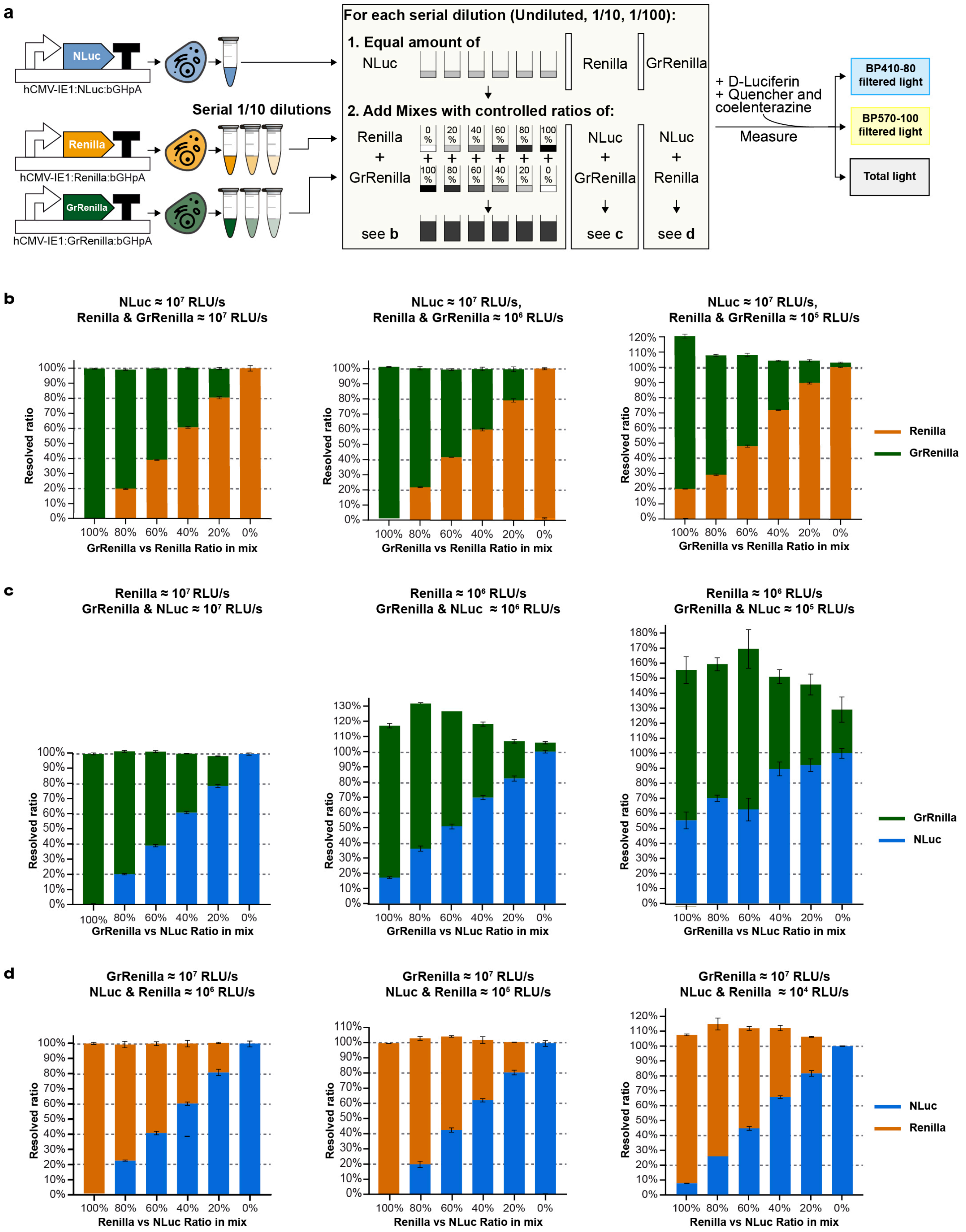
a
hCMV-IE1:NLuc:bGHpA
hCMV-IE1:Renilla:bGHpA
hCMV-IE1:GrRenilla:bGHpA
Serial 1/10 dilutions
For each serial dilution (Undiluted, 1/10, 1/100):
1. Equal amount of
NLuc
Renilla
GrRenilla
2. Add Mixes with controlled ratios of:
Renilla + GrRenilla
NLuc + GrRenilla
NLuc + Renilla
see b
see c
see d
+ D-Luciferin
+ Quencher and coelenterazine
Measure
BP410-80 filtered light
BP570-100 filtered light
Total light
b
NLuc ≈ 10⁷ RLU/s
Renilla & GrRenilla ≈ 10⁷ RLU/s
NLuc ≈ 10⁷ RLU/s,
Renilla & GrRenilla ≈ 10⁶ RLU/s
NLuc ≈ 10⁷ RLU/s,
Renilla & GrRenilla ≈ 10⁵ RLU/s
Resolved ratio
GrRenilla vs Renilla Ratio in mix
Renilla
GrRenilla
c
Renilla ≈ 10⁷ RLU/s
GrRenilla & NLuc ≈ 10⁷ RLU/s
Renilla ≈ 10⁶ RLU/s
GrRenilla & NLuc ≈ 10⁶ RLU/s
Renilla ≈ 10⁶ RLU/s
GrRenilla & NLuc ≈ 10⁵ RLU/s
Resolved ratio
GrRenilla vs NLuc Ratio in mix
GrRnilla
NLuc
d
GrRenilla ≈ 10⁷ RLU/s
NLuc & Renilla ≈ 10⁶ RLU/s
GrRenilla ≈ 10⁷ RLU/s
NLuc & Renilla ≈ 10⁵ RLU/s
GrRenilla ≈ 10⁷ RLU/s
NLuc & Renilla ≈ 10⁴ RLU/s
Resolved ratio
Renilla vs NLuc Ratio in mix
NLuc
Renilla

**Supplementary Figure 11. High expression levels of coelenterazine-responsive luciferases interfere with resolving emission signals in luciferase mixtures.** (**a**) Schematic of the experimental setup used to determine potential challenges associated with resolving emission signals in luciferase mixtures when three coelenterazine-responsive luciferases are employed. Individual plasmids, each possessing one transcriptional coelenterazine-responsive luciferase unit containing the hCMV-IE1 promoter, coelenterazine-responsive luciferase (NLuc, Renilla, or GrRenilla), and transcriptional terminator were transfected into HEK293T/17 cells. After 24 hours, transfected cells were harvested, lysed, and serial 1:10 dilutions were prepared. A defined amount of the NLuc (~$10^7$ RLU/s) normalization control was supplemented with defined amounts of Renilla and GrRenilla mixed at different ratios totaling 100% (shown). After the addition of D-Luciferin substrate-containing buffer (LARII buffer), as well as quencher and coelenterazine substrate-containing buffer (Stop & Glo buffer), total and filtered light were measured after 7 seconds (**Figure 1h**). Similar experimental setups were used to evaluate potential issues with resolving the emission signals in luciferase mixes using Renilla and GrRenilla. (**b**) Resolving the emission signals of NLuc, Renilla, and GrRenilla in a mixture works well when the brightness of all luciferases are within the same order of magnitude in brightness, i.e., levels of NLuc (~$10^7$ RLU/s) are comparable to the combined level of Renilla and GrRenilla (~$10^7$ RLU/s total levels of Renilla and GrRenilla combined) (**Left**), an order of magnitude in brightness different, i.e., levels of NLuc (~$10^7$ RLU/s) are 10x higher than the combined levels of Renilla and GrRenilla (~$10^6$ RLU/s total levels of Renilla and GrRenilla combined) (**Middle**), but not when the brightness differs by two orders of magnitude or greater, i.e., levels of NLuc (~$10^7$ RLU/s) are 100x higher than the combined levels of Renilla and GrRenilla (~$10^5$ RLU/s total levels of Renilla and GrRenilla combined) (**Right** and **data not shown**). Since NLuc is the strongest coelenterazine-responsive luciferase, spillover of its emission signals into the Renilla and GrRenilla channels results in an inaccurate contribution of the individual luciferases to the measured signal. In conclusion, use of NLuc transcriptionally driven by a strong constitutive promoter (hCMV-IE1 promoter) is not recommended for use as a control for normalization. (**c-d**) Similar observations were noted for Renilla (**c**) and GrRenilla (**d**). The separation of emission signals was problematic when the levels of Renilla were just 1 log (10x) higher compared to the combined levels of NLuc and GrRenilla (**c**, **Middle**). Four technical replicates are included in each data point, and the standard error of the mean is represented. Source data are provided as a Source Data file.

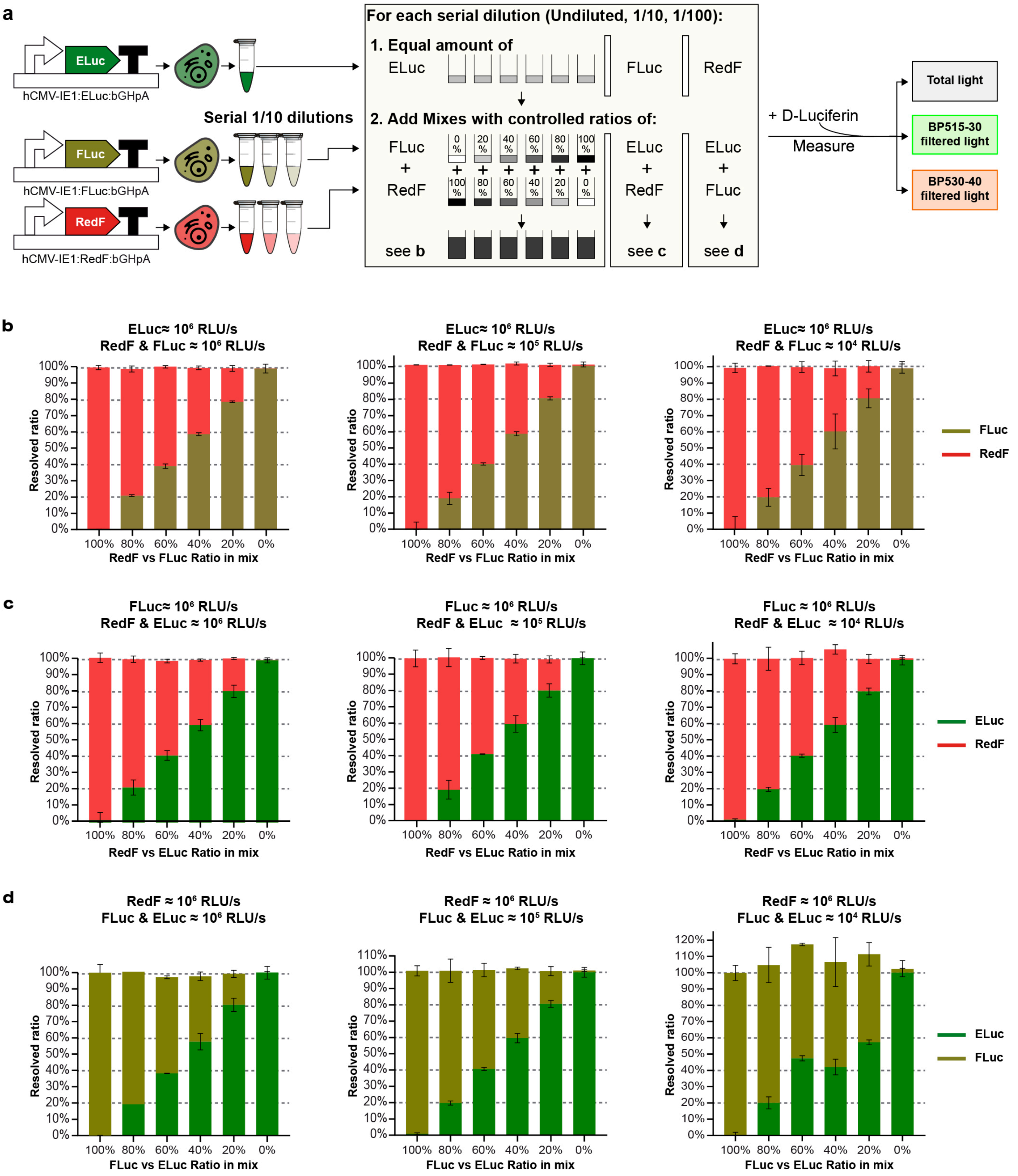


**Supplementary Figure 12. High expression levels of ELuc does not interfere with resolving RedF and ELuc emission signals in luciferase mixtures.** (**a**) Schematic of the experimental setup used to determine potential challenges with resolving the emission signals in luciferase mixtures when three D-Luciferin-responsive luciferases are employed. Individual plasmids, each possessing one transcriptional D-Luciferin-responsive luciferase unit containing the hCMV-IE1 promoter, D-Luciferin-responsive luciferase (ELuc, FLuc, or RedF), and transcriptional terminator were transfected into HEK293T/17 cells. After 24 hours, transfected cells were harvested, lysed, and serial 1:10 dilutions were prepared. To determine potential issues with resolving the emission signals in luciferase mixtures using ELuc as the normalization control, a defined amount of ELuc (~$10^6$ RLU/s) was supplemented with defined amounts of FLuc and RedF mixed at different ratios totaling 100% (shown). After the addition of D-Luciferin substrate-containing buffer (LARII buffer), total and filtered light were measured after 30 seconds (**Figure 1g**). Similar experimental setups were used to evaluate potential challenges with resolving the emission signals in luciferase mixes using FLuc and RedF. (**b**) Resolving the emission signals of ELuc, FLuc, and RedF in a mixture works well when the brightness of all luciferases are within the same order of magnitude in brightness, i.e., levels of ELuc (~$10^6$ RLU/s) are comparable to the combined levels of FLuc and RedF (~$10^6$ RLU/s total levels of FLuc and RedF combined) (**Left**), one order of magnitude in different, i.e., levels of ELuc (~$10^6$ RLU/s) are 10x higher than the combined levels of FLuc and RedF (~$10^5$ RLU/s total levels of FLuc and RedF combined) (**Middle**), and two orders of magnitude in different, i.e., levels of ELuc (~$10^6$ RLU/s) are 100x higher than the combined levels of FLuc and RedF (~$10^4$ RLU/s total levels of FLuc and RedF combined) (**Right**). In conclusion, ELuc transcriptionally driven by a strong constitutive promoter (hCMV-IE1 promoter) is a good candidate to be used as a control for normalization purposes. (**c-d**) In contrast, difficulty in resolving the emission signals of FLuc (**c**) and RedF (**d**) was observed, similar to what was observed for GrRenilla (**Supplementary Figure 11d**). In both cases, spillover of the emission signal resulted in an inaccurate contribution of the individual luciferases into the measured signal when the difference in brightness was two orders of magnitude between FLuc (~$10^6$ RLU/s) and ELuc/RedF (~$10^4$ RLU/s total levels of FLuc and RedF combined), and (**c**) RedF (~$10^6$ RLU/s) and FLuc/RedF (~$10^4$ RLU/s total levels of FLuc and RedF combined) (**d**). This analysis demonstrates that ELuc is the only luciferase in this set of six luciferases examined that can be used as a good control for normalization in the multiplex hextuple luciferase assay. Four technical replicates are included in each data point, and the standard error of the mean is represented. Source data are provided as a Source Data file.

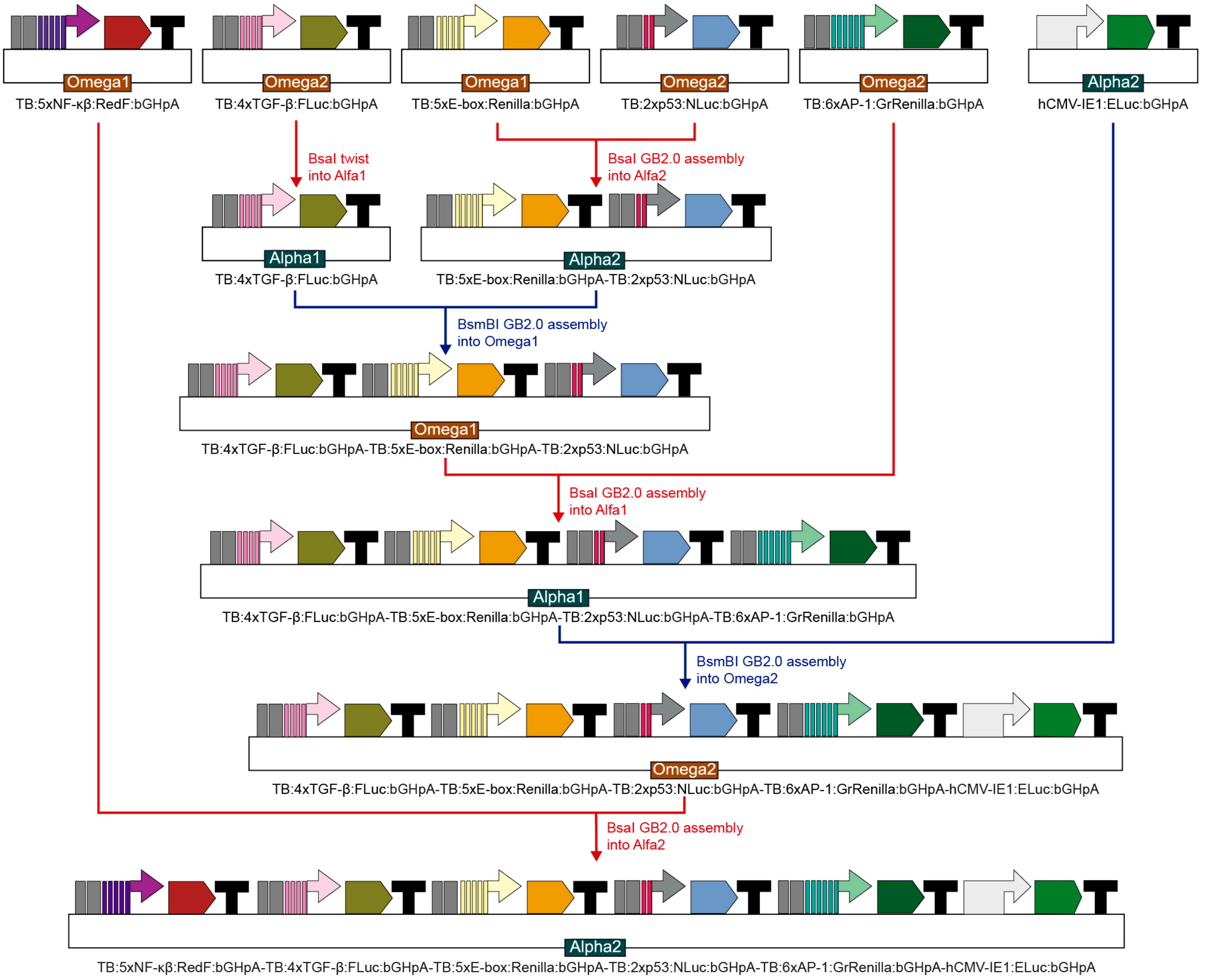


**Supplementary Figure 13. Overview of the synthetic assembly method performed to create the multi-luciferase reporter plasmid containing six luciferase transcriptional units.** In the first step, each of the transcriptional units (e.g., TB:5xNF-κβ:RedF:bGHPA) were built as described in **Supplementary Figure 10**. Next, all units were braided together in five serial steps of BsaI (red) and BsmBI (blue) assembly (**Figure 3f**). All assembly reactions were performed as described in the **Online Methods**.

**[1]** TB:5xNF-κβ:RedF:bGHpA - 4384 bp

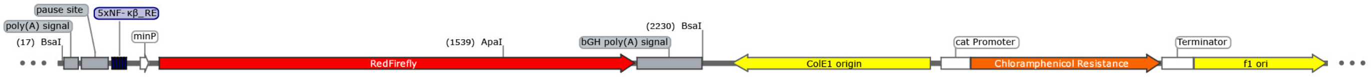


**[2]** TB:4xTGF-β:FLuc:bGHpA - 4388 bp

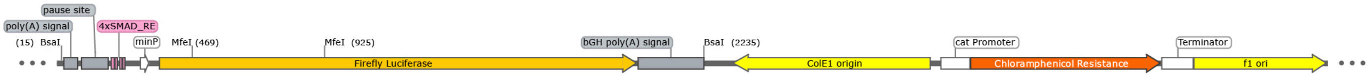


**[3]** TB:5xE-box:Renilla:bGHpA - 3656 bp

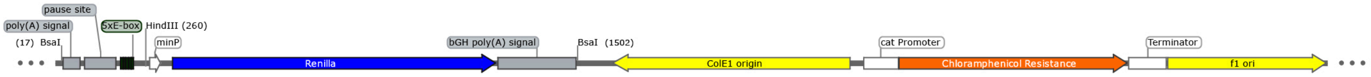


**[4]** TB:2xp53:NLuc:bGHpA - 3252 bp

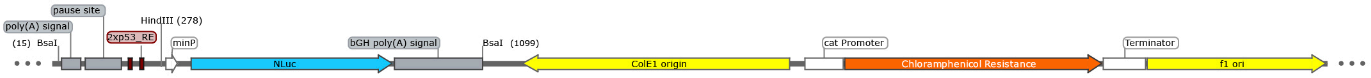


**[5]** TB:6xAP-1:GrRenilla:bGHpA -3673 bp

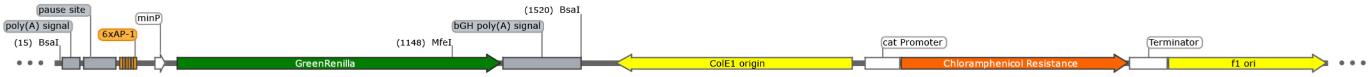


**[6]** hCMV-IE1:ELuc:bGHpA - 4844 bp

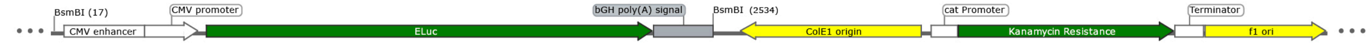


**[7]** TB:5xE-box:Renilla:bGHpA-TB:2xp53:NLuc:bGHpA - 4900 bp

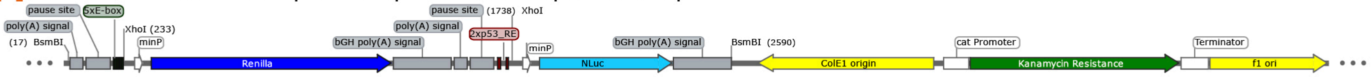


**[8]** TB:4xTGF-β:FLuc:bGHpA-TB:5xE-box:Renilla:bGHpA-TB:2xp53:NLuc:bGHpA - 6980 bp

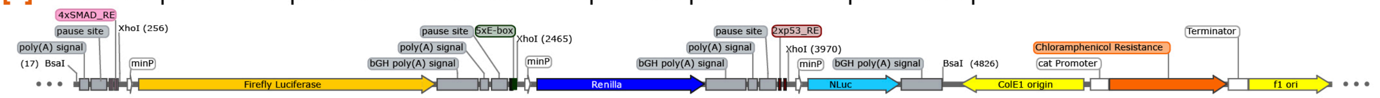


**[9]** TB:4xTGF-β:FLuc:bGHpA-TB:5xE-box:Renilla:bGHpA-TB:2xp53:NLuc:bGHpA-TB:6xAP-1:GrRenilla:bGHpA - 8644 bp

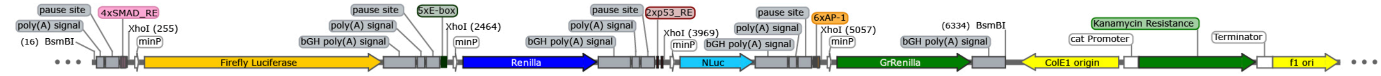


**[10]** TB:4xTGF-β:FLuc:bGHpA-TB:5xE-box:Renilla:bGHpA-TB:2xp53:NLuc:bGHpA-TB:6xAP-1:GrRenilla:bGHpA-hCMV-IE1:ELuc:bGHpA - 11007 bp

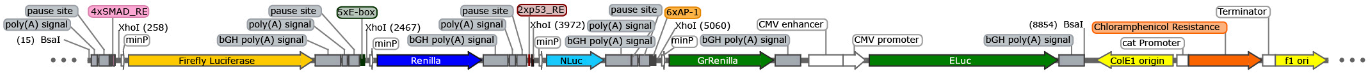


**[11]** TB:5xNF-κβ:RedF:bGHpA-TB:4xTGF-β:FLuc:bGHpA-TB:5xE-box:Renilla:bGHpA-TB:2xp53:NLuc:bGHpA-TB:6xAP-1:GrRenilla:bGHpA-hCMV-IE1:ELuc:bGHpA - 13383 bp

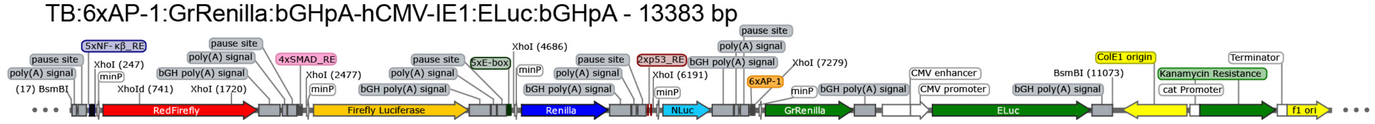

**Supplementary Figure 14. Vector maps of luciferase reporter plasmids generated in this study.** Plasmid maps of all six individual luciferase transcriptional units (Plasmids 1 to 6), intermediate assemblies (Plasmids 7 to 10), and the final hextuple luciferase vector (Plasmid 11) are shown. Important plasmid features and key restriction enzymes used for DNA fingerprinting are indicated. DNA analysis (restriction enzyme fingerprinting and uncut) of each plasmid is indicated in **Figure 3g**.

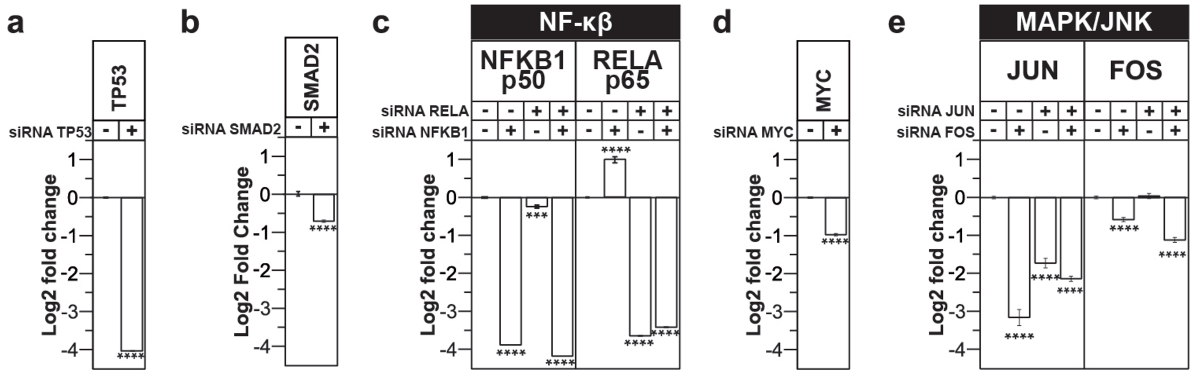


**Supplementary Figure 15. Verification of mRNA knockdown by target-specific siRNAs using quantitative PCR.** The siRNAs used were previously validated in other studies and are referenced in **Supplementary Table 6**. The primers used for quantitative PCR are listed in **Supplementary Table 10**. Effective mRNA knockdown by gene-specific siRNAs up to 16-fold was detected for all pathways: p53 (**a**), TGF-β (**b**), NF-κβ (**c**), c-Myc (**d**), and MAPK/JNK (**e**). Note: Significant upregulation of *RELA* mRNA following *NFKB1* silencing (**c**) has been reported previously[1]. Statistical significance was determined by the multiple t-test using the Holm-Sidak method with alpha = 0.05 (*P <0.05, **P < 0.01, ***P < 0.001, and ****P < 0.0001). n = 4 for all qPCR experiments. Source data are provided as a Source Data file.

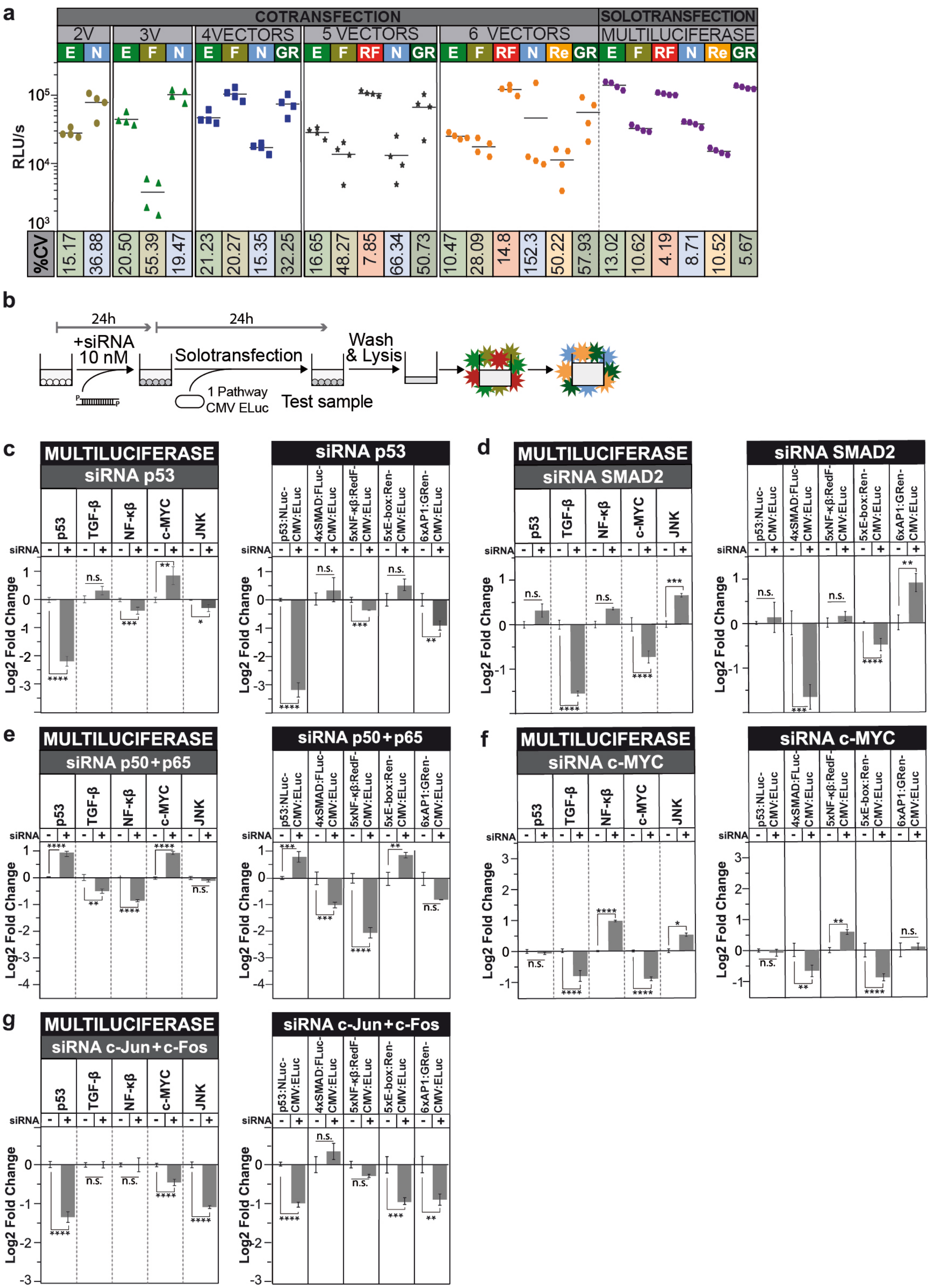


**Supplementary Figure 16. Comparison of monitoring pathway activities using the multiplex hextuple luciferase vector reporting on five pathways, and five multiplex dual luciferase vectors each reporting on just one pathway.** (**a**) Variability in the quantification of the different luciferase activities following cotransfection of two, three, four, five or six vectors, each encompassing a single luciferase reporter, and the solotransfection of all six luciferase reporters in one vector. The absolute luminescence measured as relative luminescence units per second (RLU/s) of four biological replicates, is represented on the y-axis, while the coefficient of variation (%CV) between replicates is indicated on the x-axis. A lower %CV was observed during solotransfection of all luciferase reporters incorporated in one plasmid, compared to any of the cotransfection conditions. This observation indicates that the overall error of multiplex experimentation of this kind will be lower using solotransfection of the multiplex reporter (multiluciferase) than when cotransfecting the individual plasmids encoding a combination of the single luciferase reporters ELuc (E), FLuc (F), RedF (RF), NLuc (N), Renilla (Re), and GrRenilla (GR). Four technical replicates are included in each data point; the mean is represented with the horizontal bar. (**b**) A549 cells were treated with 10 nM siRNA and incubated for 24 hours before solotransfection of the multiplex hextuple luciferase reporter vector (as shown previously in **Figure 5**), or multiplex dual luciferase plasmids that include one pathway reporter and the normalizer ELuc luciferase (this figure). After another 24 hours, cells were lysed and then multiplex hextuple luciferase assaying was performed. (**c**) The effects of siRNA silencing of *TP53* on five pathways detected by the multiplex hextuple luciferase vector (left, as shown previously in **Figure 5b**), correlate with the measurements obtained by the five multiplex dual luciferase plasmids, each reporting on one pathway (right). (**d**) The effects of siRNA knockdown of *SMAD2* on five pathways, in A459 cells previously stimulated with TGF-β, are similar when the activities are measured at once using the multiplex hextuple luciferase reporter (left, as shown previously in **Figure 5c**) or when the activities are measured separately using the five multiplex dual luciferase reporters (right). (**e**) Downregulation of the NF-κβ pathway through the simultaneous addition of siRNAs targeting *p65/RELA* and *p50/NFKB1* show similar results when the activity of the five pathways are measured using the multiplex hextuple luciferase reporter vector (left, as shown previously in **Figure 5d**) or using five multiplex dual luciferase reporter plasmids that each report on one pathway at a time (right). (**f**) The siRNA knockdown of *c-Myc/MYC* reports on-target and collateral effects (left, as shown previously in **Figure 5e**) that are corroborated when they are monitored with the five plasmids that each report on individual pathways (right). The sole exception in this case is the MAPK/JNK pathway that shows no significant change when measured in isolation (right). (**g**) The simultaneous knockdown of *c-Jun/JUN* and *c-Fos/FOS* results in decreased MAPK/JNK pathway signaling, as well as the p53 and c-Myc pathways. These results were independently obtained using the multiplex hextuple luciferase vector reporting on all five pathways at once (left, as shown previously in

**Figure 5f**) and the five multiplex dual luciferase plasmids that each report on a single pathway (right). Source data are provided as a Source Data file.

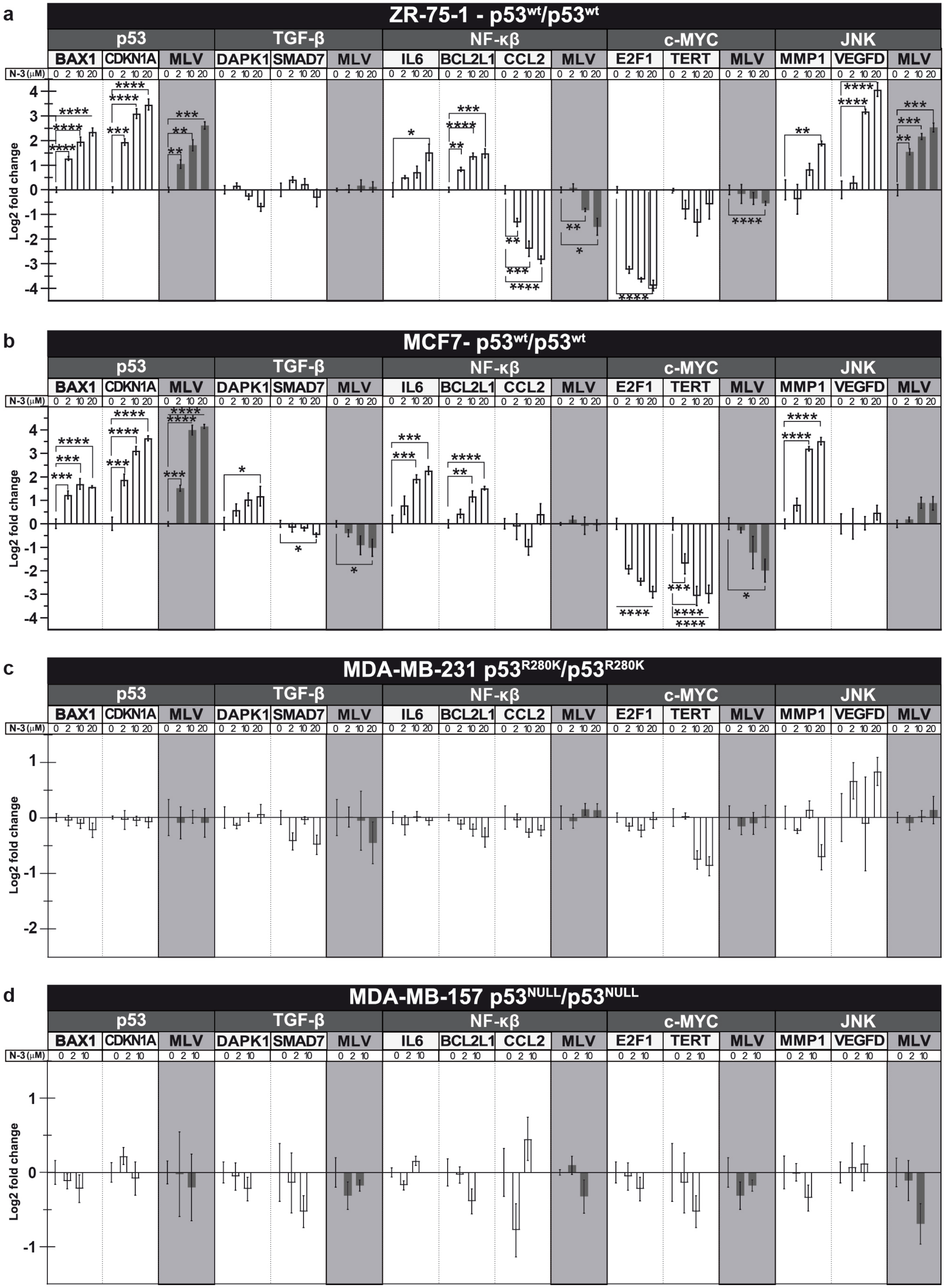
a
ZR-75-1 - p53wt/p53wt
b
MCF7- p53wt/p53wt
c
MDA-MB-231 p53R280K/p53R280K
d
MDA-MB-157 p53NULL/p53NULL
p53
TGF-β
NF-κβ
c-MYC
JNK
BAX1
CDKN1A
MLV
DAPK1
SMAD7
IL6
BCL2L1
CCL2
E2F1
TERT
MMP1
VEGFD
N-3 (μM)
Log2 fold change

**Supplementary Figure 17. Collateral effects of Nutlin-3 treatment are detected by the multiplex luciferase assay and qPCR.** (**a-b**) Activation of the p53 pathway was observed in the two Nutlin-3-sensitive $TP53^{WT}$ cell lines, ZR-75-1 (**a**) and MCF-7 (**b**), by qPCR and the multiplex luciferase assay. Most collateral effects were correlated between qPCR and luciferase emission recordings. The only exception was the NF-κβ pathway. Activation of two downstream genes (*IL6* and *BLC2L1*) was detected by qPCR. However, luciferase measurements produced results indicating downregulation and no significant effect in ZR-75-1 (**a**) and MCF7 (**b**) cell lines, respectively. Previous studies reported that Nutlin-3 inhibits the NF-κβ pathway in a p53-dependent manner in A459 cells[2] or imparts no effect on IL6 expression in MCF-7 cells[3], suggesting that the effects of Nutlin-3 are context-dependent (**c-d**). No significant changes were detected for the MDA-MB-231 ($TP53^{R280K}$) (**c**) and MDA-MV-157 ($TP53^{NULL}$) (**d**) cell lines by qPCR or the multiplex luciferase assay. Statistical significance was determined by the multiple t-test using the Holm-Sidak method with alpha = 0.05 (*P <0.05, **P < 0.01, ***P < 0.001, and ****P < 0.0001). n = 4 for all qPCR experiments. Source data are provided as a Source Data file.

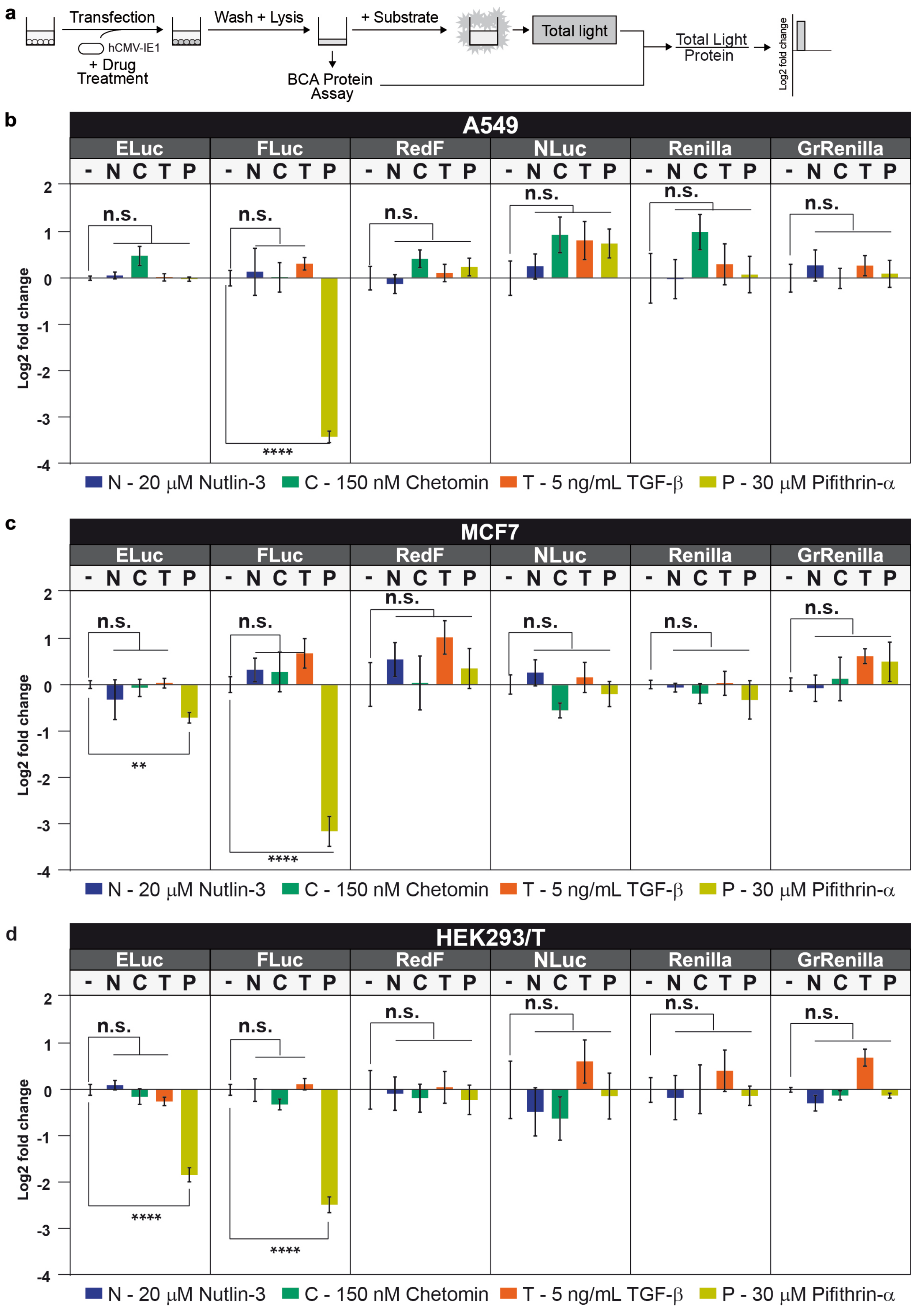
a
Transfection
hCMV-IE1
+ Drug Treatment
Wash + Lysis
+ Substrate
Total light
BCA Protein Assay
Total Light
Protein
Log2 fold change
b
A549
ELuc
FLuc
RedF
NLuc
Renilla
GrRenilla
- N C T P
n.s.
****
Log2 fold change
N - 20 μM Nutlin-3
C - 150 nM Chetomin
T - 5 ng/mL TGF-β
P - 30 μM Pifithrin-α
c
MCF7
**
d
HEK293/T

**Supplementary Figure 18. Luciferase activities are not influenced by the drugs used in this study.** (**a**) Schematic of the experimental setup used to determine luciferase activity interference. To exclude the possibility that drug treatments may inhibit one or more of the luciferase activities, drugs were assayed against each constitutively expressed luciferase. Cells were transfected with a luciferase reporter plasmid, encoding one of the six luciferases (ELuc, FLuc, RedF, NLuc, Renilla, and GrRenilla) driven by the constitutive hCMV-IE1 promoter, and treated with or without the highest concentration of the drugs used during this study: 20 μM Nutlin-3 (N), 150 nM Chetomin (C), and 5 ng/ml TGF-β (T). After cell lysis, total protein content was measured by the BCA protein assay, and total light emitted by each luciferase was measured after addition of appropriate substrate-containing buffer, *i.e.*, D-Luciferin substrate-containing buffer (LARII buffer) for the D-Luciferin luciferases, or quencher and coelenterazine substrate-containing buffer (Stop & Glo buffer) for the coelenterazine luciferases. Values obtained from drug-treated cells were normalized against values obtained from non-treated cells. As a positive control, cells were treated with 30 μM Pifithrin-α (P), which at that concentration is a well-known *in vitro* and *in vivo* inhibitor of the activity of the firefly luciferase, FLuc[4]. (**b-d**) None of the drugs used in this work have a significant off-target effect on the light emitted by the constitutively expressed luciferases using A549 cells (b), and MCF-7 cells (c). On the other hand, Pifithrin-α treatment resulted in a 3.4 (**b**) and 3.2 (**c**), and 2.47 (**d**) fold reduction of the light emitted by FLuc, as expected. Also, it resulted in a 0.71 (**c**) and 1.83 (**d**) fold reduction of the light emitted by ELuc, which has a similar bioluminescent mechanism to FLuc. Statistical significance was determined by the multiple t-test using the Holm-Sidak method with alpha = 0.05 (*P <0.05, **P < 0.01, ***P < 0.001, and ****P < 0.0001). n = 4 for all experiments in luminescence measurement, and the protein content of each well was calculated in with three technical replicates in a BCA assay. Source data are provided as a Source Data file.

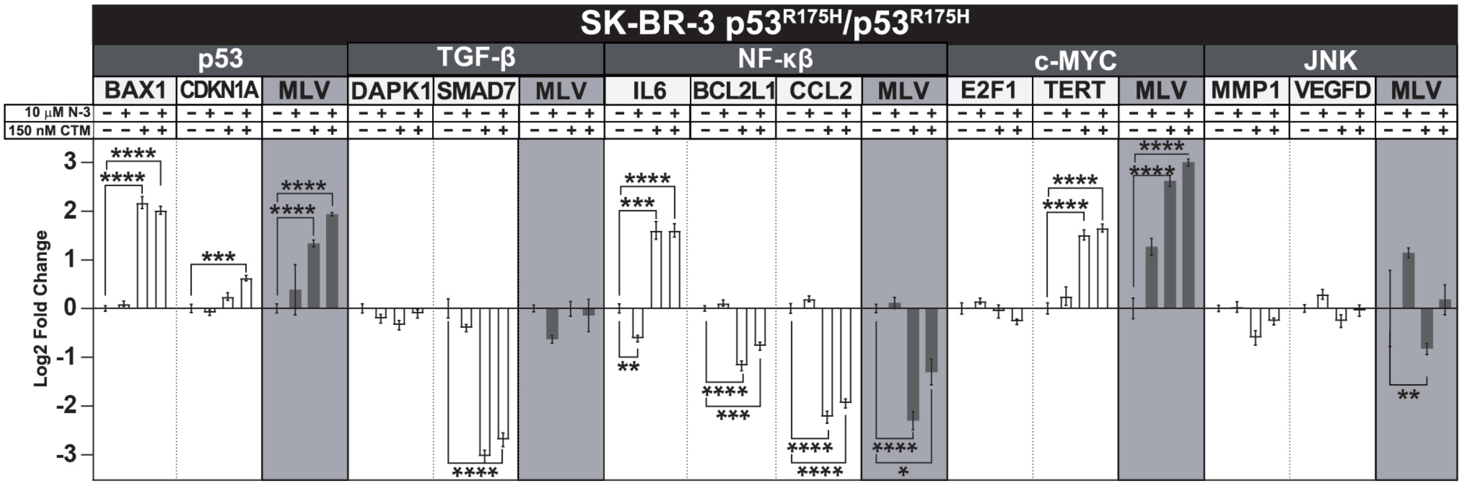


**Supplementary Figure 19. Collateral effects of Chetomin treatment of the SK-BR-3 cell line ($TP53^{R175H}$) are detected by the multiplex luciferase assay and qPCR.** The correlation between qPCR results and luciferase emission recordings, obtained after transfection of the multiplex luciferase vector (MLV), demonstrate the complementary effects of Nutlin-3 and Chetomin treatment on p53 signaling. While $TP53^{R175H}$ is not significantly activated with Nutlin-3 alone, the addition of Chetomin strongly reactivates this pathway. This level of activation is even further enhanced in the combined presence of Chetomin and Nutlin-3. Most of the collateral and differential effects observed with luciferase emission recordings on the other pathways were also corroborated by qPCR analysis. Statistical significance was determined by the multiple t-test using the Holm-Sidak method with alpha = 0.05 (*P <0.05, **P < 0.01, ***P < 0.001, and ****P < 0.0001). n = 4 for all qPCR experiments. Source data are provided as a Source Data file.

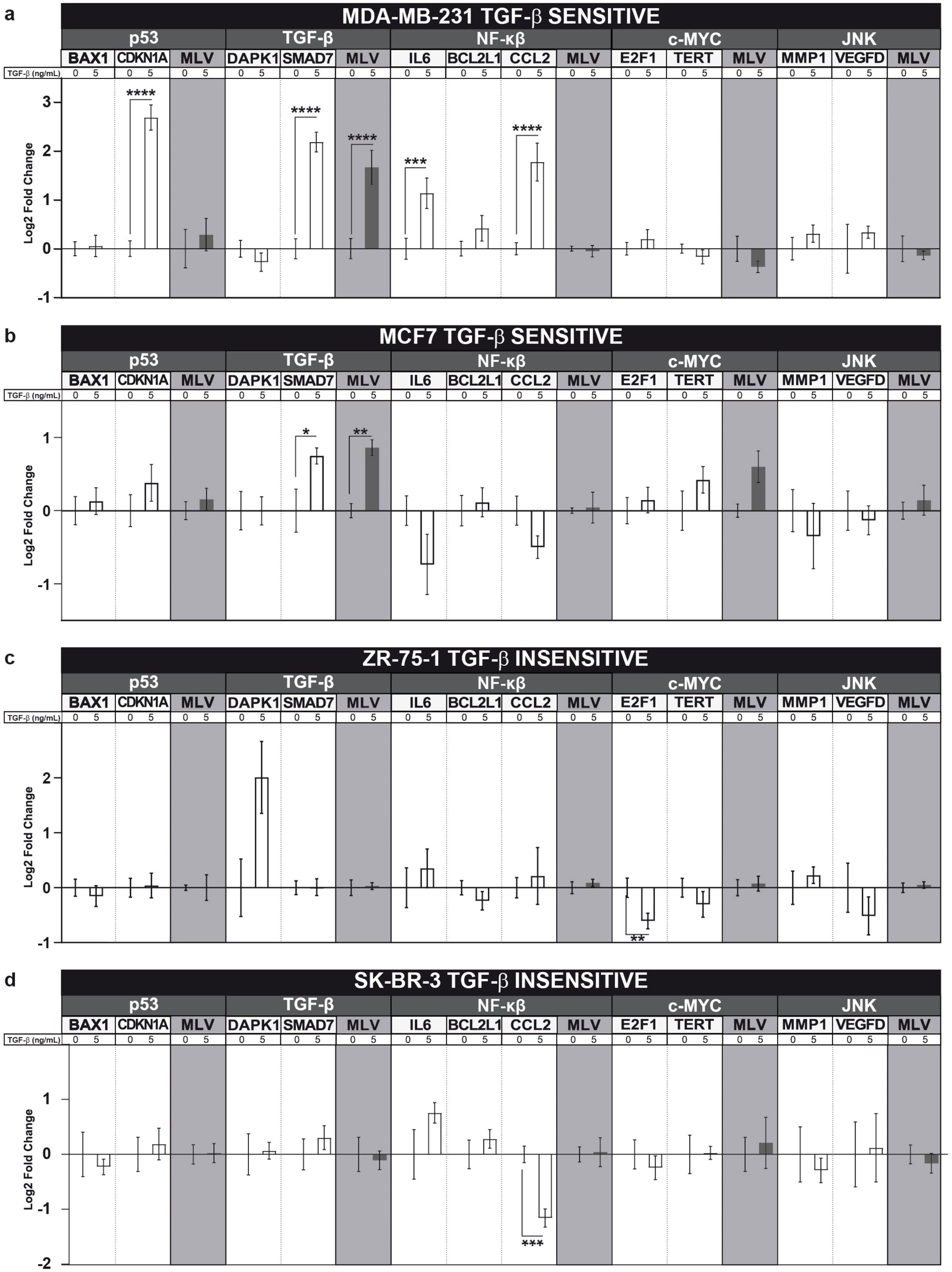
a
MDA-MB-231 TGF-β SENSITIVE
b
MCF7 TGF-β SENSITIVE
c
ZR-75-1 TGF-β INSENSITIVE
d
SK-BR-3 TGF-β INSENSITIVE
p53
TGF-β
NF-κβ
c-MYC
JNK
BAX1
CDKN1A
MLV
DAPK1
SMAD7
IL6
BCL2L1
CCL2
E2F1
TERT
MMP1
VEGFD
TGF-β (ng/mL)
Log2 Fold Change

**Supplementary Figure 20. Collateral effects of recombinant TGF-β-mediated activation of the TGF-β cellular signaling pathway are detected by the multiplex luciferase assay and qPCR.** (**a-b**) Two TGF-β–sensitive cell lines, MDA-MB-231 (**a**) and MCF7 (**b**), exhibit TGF-β pathway activation after 6-hour treatment with recombinant TGF-β. Interestingly, while no significant collateral effects were detected for the synthetic p53 and NF-κβ transcriptional reporters using the multiplex luciferase assay, significant activation of downstream gene expression (*CDKN1A* for the p53 pathway, *IL6* and *CCL2* for the NF-κβ pathway) was observed by qPCR in the MDA-MB-231 cell line (**a**). These observations are consistent with previous findings from other cell lines that demonstrated the presence of SMAD-binding elements in the promoters of *CDKN1A*[5] and *CCL2*[6] or direct crosstalk between the TGF-β and NF-κβ signaling pathways within the *IL6* promoter[7,8]. (**c-d**) TGF-β pathway activation was not observed in two TGF-β–insensitive cell lines, ZR-75-1 (**c**) and SK-BR-3 (**d**), through qPCR or the multiplex luciferase assay. Interestingly, qPCR revealed collateral effects in both cases: *E2F1* downregulation in ZR-75-1 cells (**c**) and *CCL2* downregulation in SK-BR-3 cells (**d**). Statistical significance was determined by multiple t-test using the Holm-Sidak method with alpha = 0.05 (*P <0.05, **P < 0.01, ***P < 0.001, and ****P < 0.0001). n = 4 for all qPCR experiments. Source data are provided as a Source Data file.

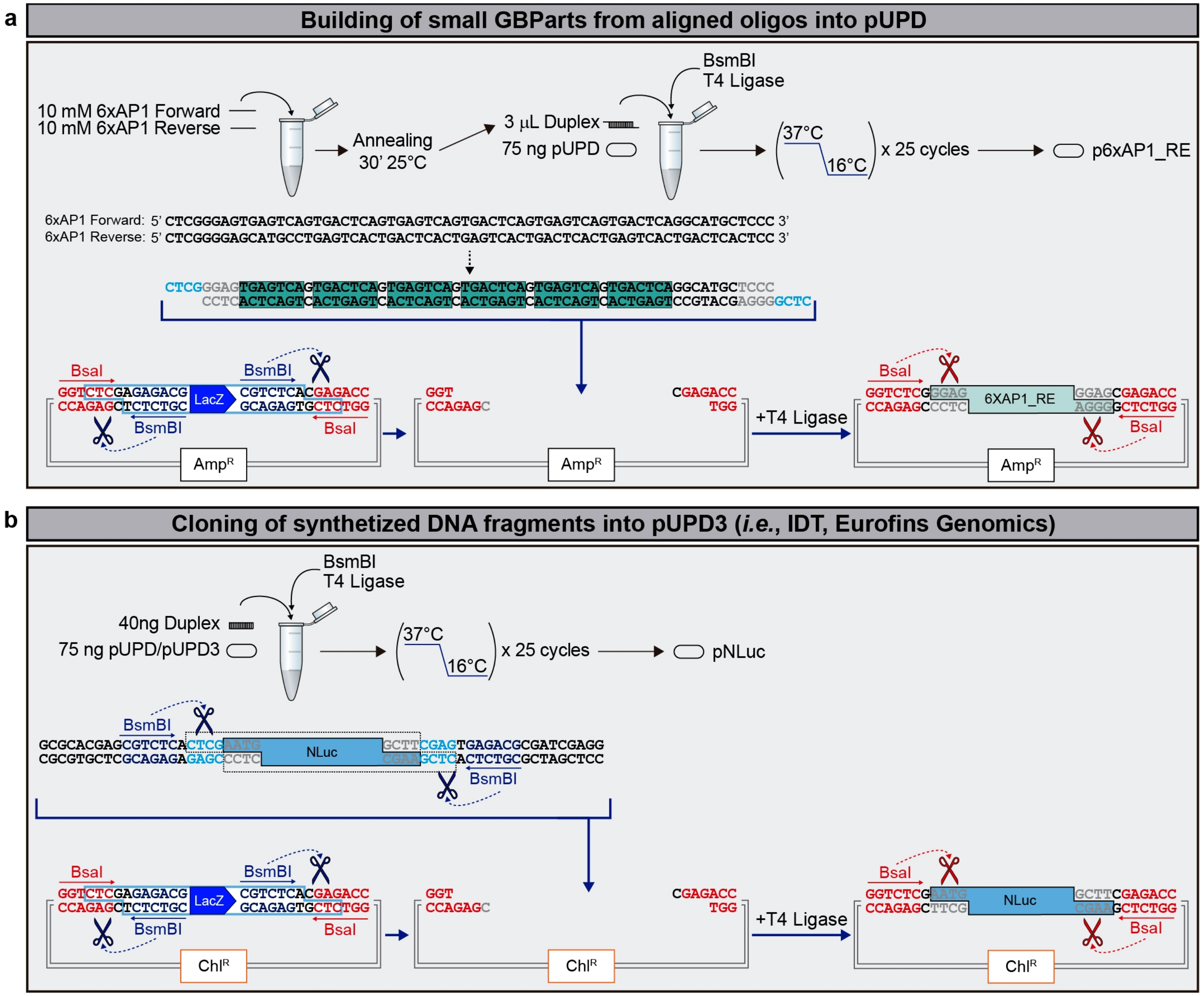


**Supplementary Figure 21. Adaptation, or "domestication", of synthetic DNA fragments in the synthetic assembly pipeline.** (**a**) Small synthetic DNA fragments (*e.g.*, 6xAP1_RE) were built from two annealed oligonucleotides in a one-step, one-pot GoldenBraid 2.0 assembly reaction into the pUPD vector[9] using BsmBI and T4 ligase. Briefly, 10 mM of both oligonucleotides were annealed for 30 minutes at 25°C. To generate 6xAP1_RE GBPart, 3 µl of the annealing reaction was combined with 75 ng of pUPD destination vector, T4 ligase, and buffer, as previously described for generating GBParts from oligo duplexes[10]. (**b**) Double-stranded synthetic DNA fragments (*e.g.*, NLuc) were cloned into the pUPD3 vector in a one-step, one-pot GoldenBraid 2.0 assembly reaction using BsmBI and T4 ligase. Briefly, 40 ng of synthetized fragment was combined with 75 ng of pUPD3 destination vector, T4 ligase, and buffer as established by GoldenBraid 2.0 rules. A similar assembly was performed using the pUPD vector[9] to generate pRedF, following the same protocol but using ampicillin instead of chloramphenicol for selection. $Amp^R$ and $Chl^R$ stand for ampicillin- and chloramphenicol-resistance markers for bacterial selection of the DNA clones.

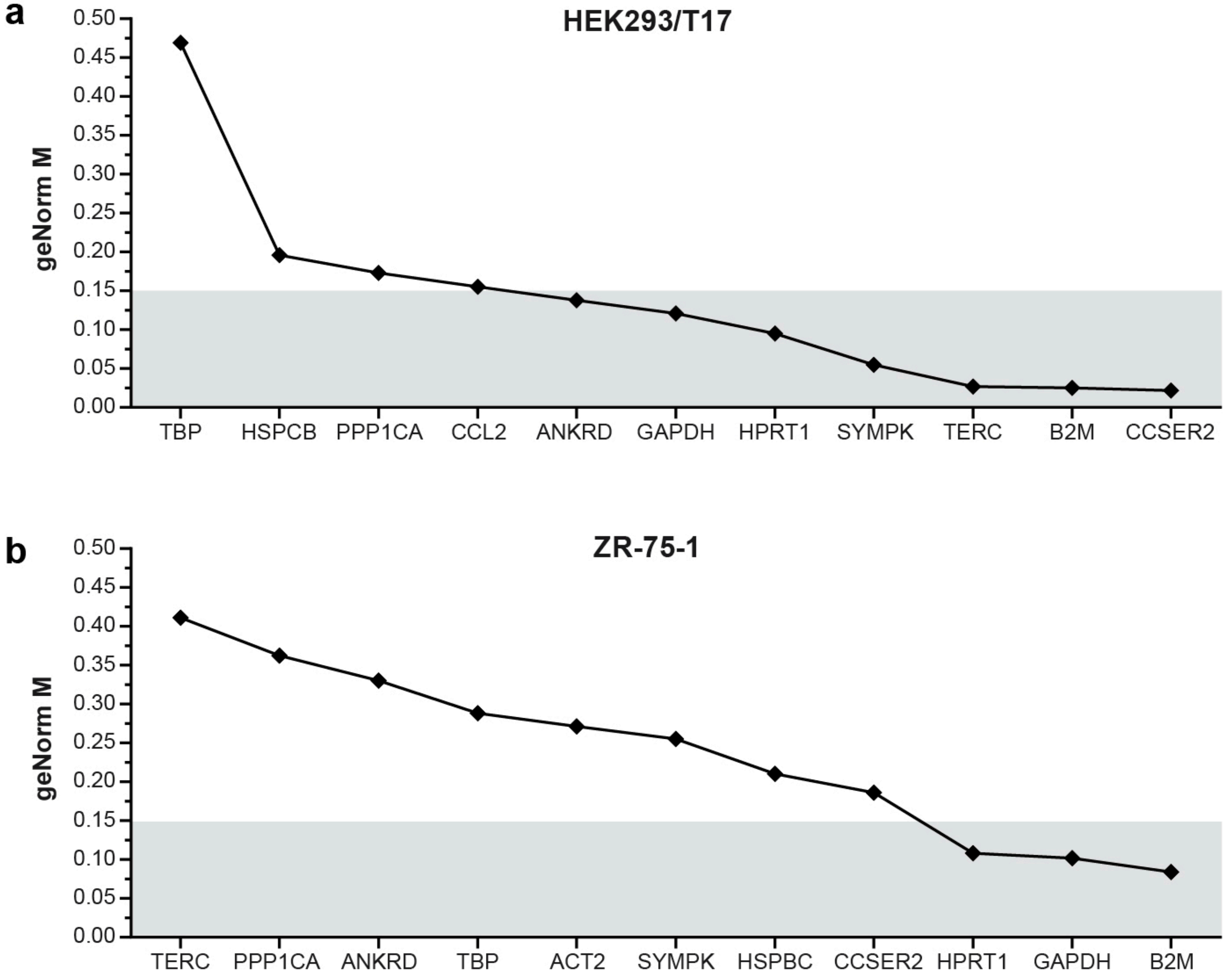


**Supplementary Figure 22. Determination of the most stable housekeeping genes for qPCR normalization.** (**a-b**) qPCR normalization was performed using the qbase+ program (Biogazelle). The optimal number of reference targets for housekeeping genes reported by the qbase+ software is 2 (geNorm V < 0.15 when comparing a normalization factor based on the two or three most stable targets, indicated by a grey area). Using this analysis, the optimal normalization factor can be calculated as the geometric mean of reference targets B2M and CCSER2 for HEK293/T17 cells (**a**) or GAPDH and B2M for ZR-75-1 cells (**b**). These values denote very high reference target stability (average geNorm M ≤ 0.2), which is typically seen when evaluating reference targets using genomic DNA as input (the quantity of any genomic reference target from the same amount of cells using the same preparation is very similar between different samples) versus

RNA (the quantity of any RNA reference target from the same amount of cells using the same preparation is dependent on a variety of factors, transcription, stability, etc.). RNA reference targets can be highly variable unless they are very stably expressed. For these reasons, the five most stable housekeeping genes (HPRT1, SYMPK, GAPDH, CCSER2, and B2M) were incorporated as reference genes in the qPCR panel and the two best performing ones were used for further normalization analysis.

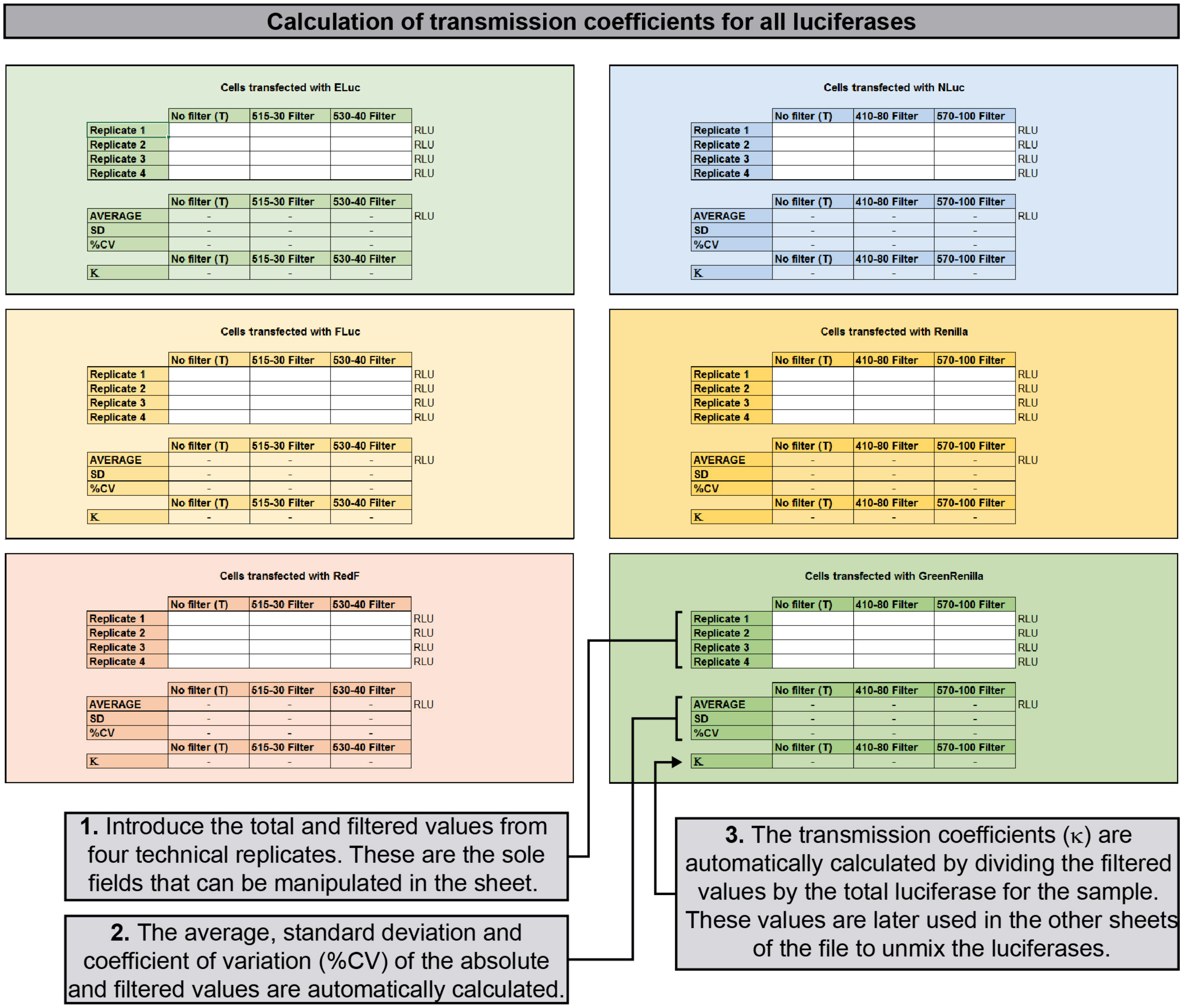


**Supplementary Figure 23. Calculation of transmission coefficients.** A screenshot of the first sheet of the provided Microsoft Excel template, including simplified instructions to facilitate calculation of the transmission coefficients for GrRenilla. Please follow the same instructions for the other five luciferases (ELuc, FLuc, RedF, NLuc, and Renilla) to calculate all necessary transmission coefficients.

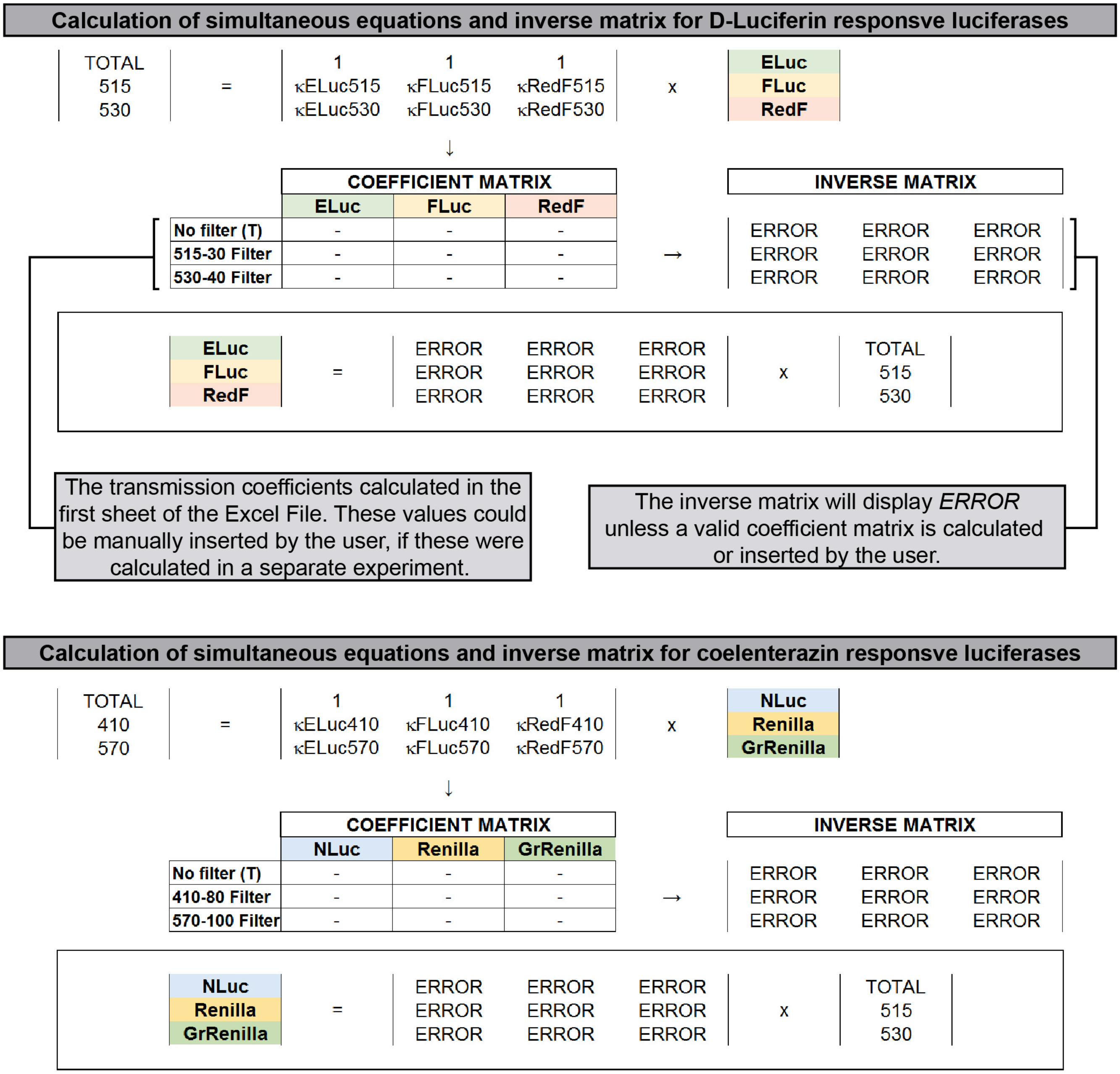


**Supplementary Figure 24. Calculation of simultaneous equations.** A screenshot of the second sheet of the provided Microsoft Excel template, including simplified instructions to facilitate generation of the simultaneous equations needed to deconvolute the obtained measurements into distinct values for all six luciferases.

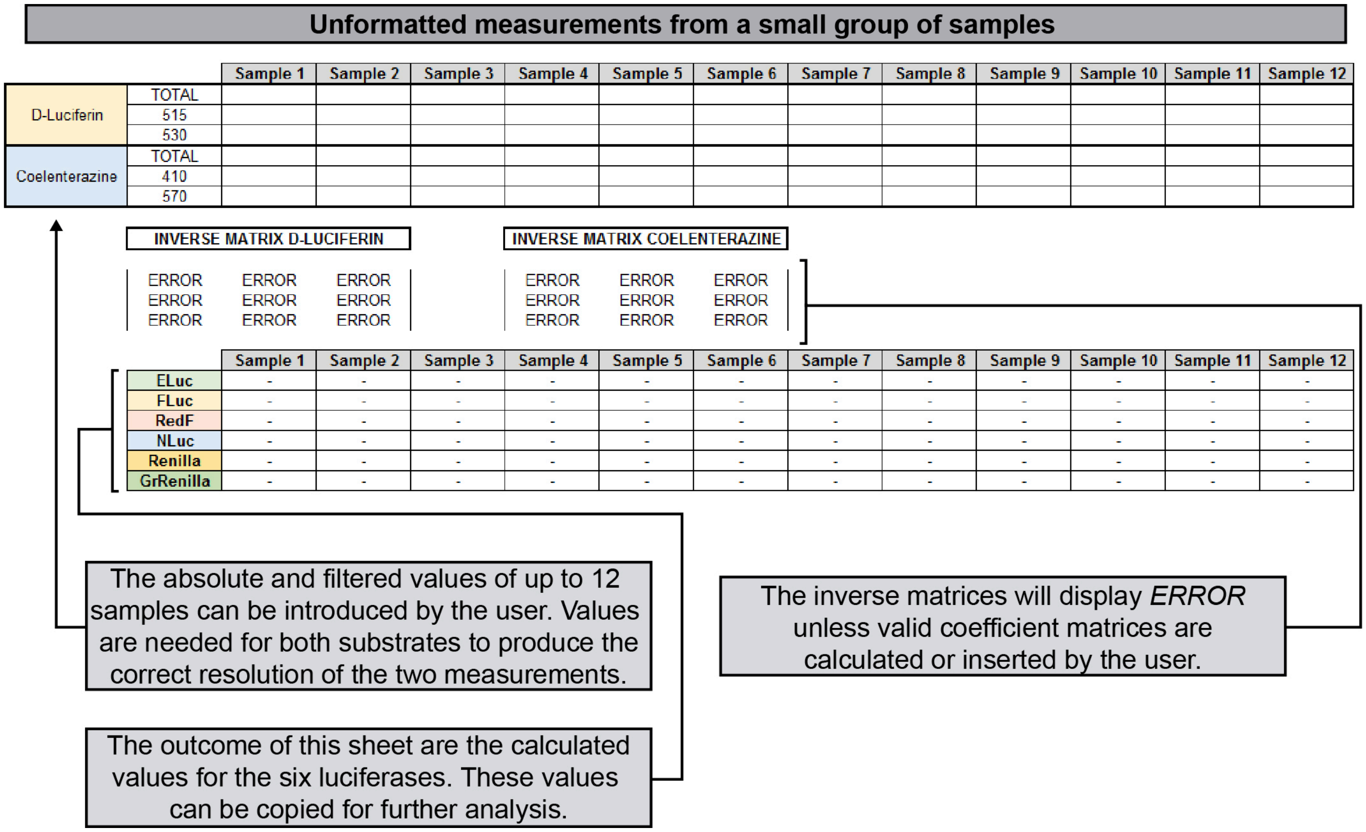


**Supplementary Figure 25. Calculation of unformatted measurements from a small group of samples.** Screenshot of the third sheet of the provided Microsoft Excel template, including simplified instructions on how to generate output values of the six luciferases from the measured values generated by the plate leader. This sheet is prepared to process up to 12 samples.

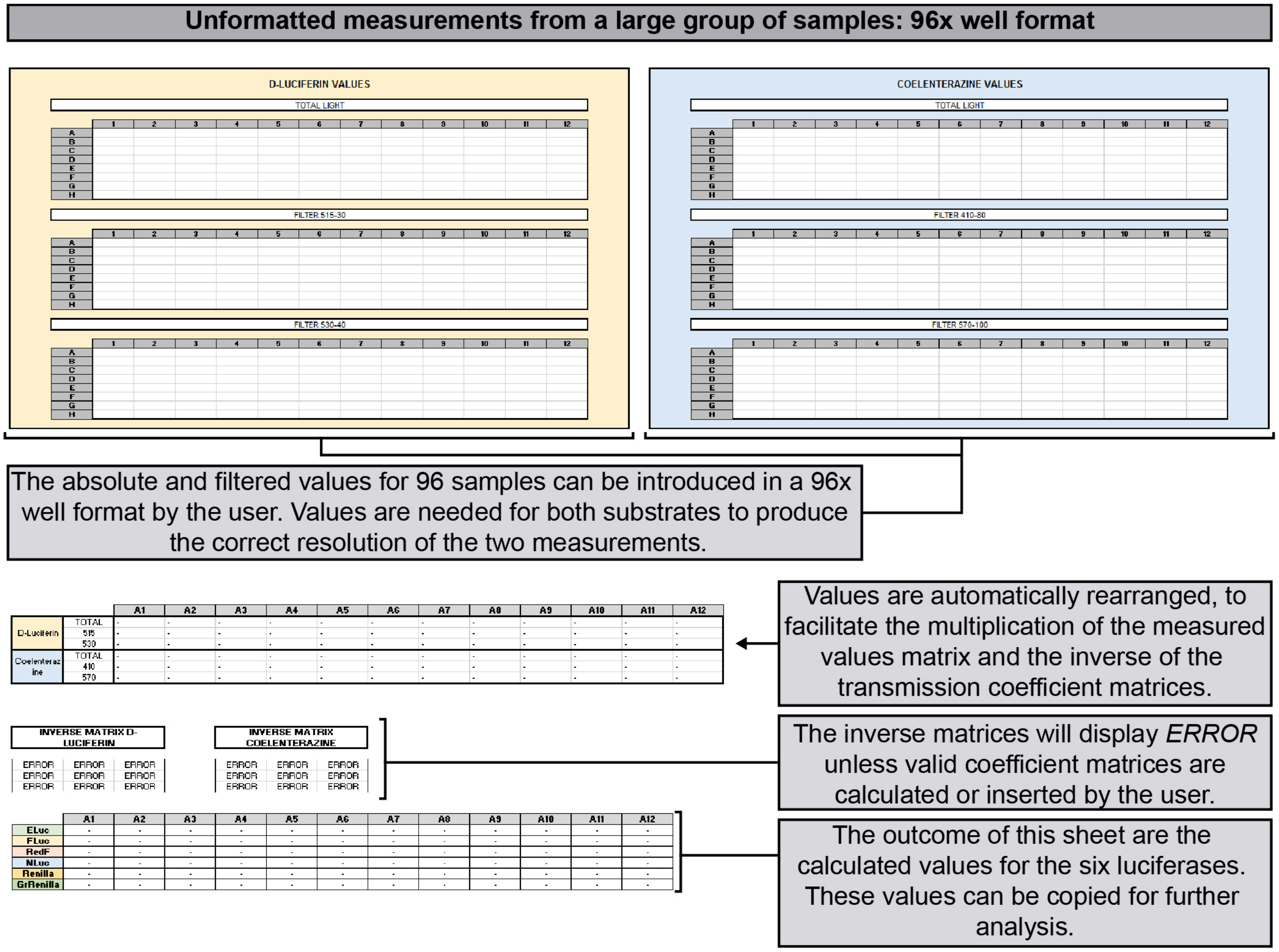


**Supplementary Figure 26. Calculation of unformatted measurements from a large group of samples.** Screenshot of portions of the fourth sheet of the provided Microsoft Excel template, including simplified instructions on how to generate output values of the six luciferases from measured values generated by the plate leader. This sheet is prepared to process samples from 96-well plates.

**Supplementary Tables**

| | ELuc | | FLuc | | RedF | |
|---|---|---|---|---|---|---|
| | **BP515-30** | **BP530-40** | **BP515-30** | **BP530-40** | **BP515-30** | **BP530-40** |
| **A549** | 24.45 ± 0.04% | 46.15 ± 0.04% | 7.24 ± 0.02% | 29.80 ± 0.04% | 0.11 ± 0.00% | 1.36 ± 0.01% |
| **MCF7** | 24.22 ± 0.28% | 46.04 ± 0.27% | 7.28 ± 0.04% | 29.89 ± 0.04% | 0.98 ± 0.00% | 1.27 ± 0.02% |
| **MDA-MB-231** | 24.32 ± 0.41% | 46.17 ± 0.50% | 7.29 ± 0.03% | 29.90 ± 0.20% | 0.10 ± 0.01% | 1.27 ± 0.020% |
| **SK-BR-3** | 24.23 ± 0.76% | 46.23 ± 1.30% | 7.29 ± 0.03% | 30.01 ± 0.53% | 0.19 ± 0.47% | 1.34 ± 0.15% |

| | NLuc | | Renilla | | GrRenilla | |
|---|---|---|---|---|---|---|
| | **BP410-80** | **BP570-100** | **BP410-80** | **BP570-100** | **BP410-80** | **BP570-100** |
| **A549** | 15.82 ± 0.18% | 11.27 ± 0.09% | 5.15 ± 0.06% | 25.52 ± 0.15% | 2.43 ± 0.04% | 53.40 ± 0.07% |
| **MCF7** | 14.20 ± 0.189% | 13.61 ± 0.18% | 4.95 ± 0.03% | 26.40 ± 0.09% | 2.13 ± 0.01% | 56.06 ± 0.05% |
| **MDA-MB-231** | 15.02 ± 0.05% | 12.14 ± 0.03% | 4.97 ± 0.04% | 26.57 ± 0.12% | 2.00 ± 0.01% | 56.69 ± 0.06% |
| **SK-BR-3** | 14.99 ± 0.123% | 12.24 ± 0.10% | 4.90 ± 0.04% | 26.56 ± 0.09% | 2.07 ± 0.04% | 56.14 ± 0.12% |

**Supplementary Table 1. Transmission coefficients of the six luciferases selected for the multiplex luciferase assay in different human cell lines**. Each luciferase was expressed in four human cell lines: A549, MCF7, MDA-MB-231, and SK-BR-3 cells (**Supplementary Table 8** for additional information related to the cell lines). Transmission coefficients were consistent between the cell lines. Statistical significance of differences observed in the transmission coefficients was determined by Tukey multiple pairwise-comparison (*P <0.05). Differences between the values calculated for the six luciferases in the four cell lines were not found to be statistically significant. Source data are provided as a Source Data file.

| | ELuc | | FLuc | | RedF | |
|---|---|---|---|---|---|---|
| **RLU** | **BP515-30** | **BP530-40** | **BP515-30** | **BP530-40** | **BP515-30** | **BP530-40** |
| **$10^7$** | 24.45 ± 0.04% | 46.15 ± 0.04% | 7.24 ± 0.02% | 29.80 ± 0.04% | 0.11 ± 0.00% | 1.36 ± 0.01% |
| **$10^6$** | 24.22 ± 0.28% | 46.04 ± 0.27% | 7.28 ± 0.04% | 29.89 ± 0.04% | 0.98 ± 0.00% | 1.27 ± 0.02% |
| **$10^5$** | 24.32 ± 0.41% | 46.17 ± 0.50% | 7.29 ± 0.03% | 29.90 ± 0.20% | 0.10 ± 0.01% | 1.27 ± 0.020% |
| **$10^4$** | 24.23 ± 0.76% | 46.23 ± 1.30% | 7.29 ± 0.03% | 30.01 ± 0.53% | 0.19 ± 0.47% | 1.34 ± 0.15% |
| **$10^3$** | 23.77 ± 3.24% | 46.66 ± 3.07%* | 8.46 ± 0.08%* | 30.07 ± 0.53%* | 0.556 ± 0.87%* | 1.78 ± 0.30%* |

| | NLuc | | Renilla | | GrRenilla | |
|---|---|---|---|---|---|---|
| **RLU** | **BP410-80** | **BP570-100** | **BP410-80** | **BP570-100** | **BP410-80** | **BP570-100** |
| **$10^7$** | 14.62 ± 0.12% | 12.88± 0.13% | 4.46 ± 0.012% | 27.93 ± 0.07% | 1.94 ± 0.27% | 57.42 ± 0.38% |
| **$10^6$** | 14.63 ± 0.08% | 12.86 ± 0.12% | 4.45 ± 0.02% | 27.96 ± 0.12% | 1.92 ± 0.02% | 57.41 ± 0.05% |
| **$10^5$** | 14.61 ± 0.08% | 12.84 ± 0.20% | 4.51 ± 0.04% | 28.11 ± 0.07% | 1.96 ± 0.05% | 57.42 ± 0.13% |
| **$10^4$** | 14.90 ± 0.40% | 16.956 ± 2.19%* | 4.44 ± 0.03% | 30.68 ± 1.80% | 2.41 ± 0.06% | 57.17 ± 1.24% |
| **$10^3$** | 11.45 ± 2.66%* | 28.54 ± 5.24%* | 4.62 ± 0.47% | 32.16± 4.77%* | 2.86 ± 1.19% | 62.33 ± 3.45%* |

**Supplementary Table 2. Dynamic luminescence range for the six luciferases selected for the multiplex luciferase assay.** Each luciferase was transfected into HEK293T/17 cells to determine their dynamic luminescence range. Lysates were diluted (1:10, 1:100, 1:1000, and so on) in Passive Lysis Buffer followed by luminescence measurements to probe the detection range using the CLARIOStar microplate reader (see **Online Methods** for hardware details). Transmission coefficients were found to be consistent from $10^7$ to $10^4$ RLU/s for both the D-Luciferin- and coelenterazine-responsive luciferases. Statistical significance of the differences observed in the transmission coefficient was determined by Tukey multiple pairwise-comparison (*P <0.05). Source data are provided as a Source Data file**.**

| Pathway | DNA binding motif | Copy number | Binding proteins | Reference |
|---|---|---|---|---|
| **p53** | RRRCWWGYYY | 2 | Two TP53 tetramers | [11] |
| | AGGCAAGTCC and AGACATGTTCT | | | |
| **TGF-β** | GTCTAGAC | 4 | SMAD2/SMAD3 heterodimer | [12] |
| **NF-κβ** | GGGPuNNPyPyCC | 5 | NFKB1/RELA heterodimer | [13] |
| | 3 x GGGAATTTCC and 2 x GGGACTTT | | | |
| **c-Myc** | CACGTG (E-Box) | 5 | MYC/MAX heterodimer | [14,15] |
| **MAPK/JNK** | TGAGTCA | 6 | JUN/FOS heterodimer | [16] |

**Supplementary Table 3. Transcriptional response elements for cellular signaling pathways used in this study.**

| Type | Abbreviation | Description | Function | Vector | Resistance | Addgene |
|---|---|---|---|---|---|---|
| Domestication | pUPD | Universal Part Domesticator | Building new basic parts | ColE1 vector | Ampicillin | [9] |
| | pUPD3 | Universal Part Domesticator #3 | Building new basic parts | ColE1 vector | Chloramphenicol | #118043 |
| Vectors | Alpha1 | pColE1_Alpha1 | Empty Vector | ColE1 vector | Kanamycin | #118044 |
| | Alpha2 | pColE1_Alpha2 | Empty Vector | ColE1 vector | Kanamycin | #118045 |
| | Omega 1 | pColE1_Omega1 | Empty Vector | ColE1 vector | Chloramphenicol | #118046 |
| | Omega 2 | pColE1_Omega2 | Empty Vector | ColE1 vector | Chloramphenicol | #118047 |
| GoldenBraid2.0 Basic Parts | phCMV-IE1 | Cytomegalovirus enhancer and promoter | High level expression constitutive promoter | pML1 | Ampicillin | #118048 |
| | pMiniP | Minimal promoter | TATA box and minimal promoter | pUPD3 | Chloramphenicol | #118049 |
| | p5xNF-κβ_RE | 5 copies of the NF-κβ DNA binding motif | Operator – DNA Response element | pUPD3 | Chloramphenicol | #118050 |
| | p4xSMAD_RE | 4 copies the SMAD DNA binding motif | Operator – DNA Response element | pUPD | Ampicillin | #118051 |
| | p5xE-Box | 5 copies of the E-box motif | Operator – DNA Response element | pUPD3 | Chloramphenicol | #118052 |
| | p2xp53_RE | 2 copies of the p53 DNA binding motif | Operator – DNA Response element | pUPD3 | Chloramphenicol | #118053 |
| | p6xAP-1_RE | 6 copies of the AP-1 binding motif | Operator – DNA Response element | pUPD | Ampicillin | #118054 |
| | pELuc | ELuc Luciferase CDS | CDS + STOP Codon | pUPD3 | Chloramphenicol | #118056 |
| | pFLuc | FLuc Luciferase | CDS + STOP Codon | pUPD | Ampicillin | #68201[9] |
| | pRedF | RedF Luciferase | CDS + STOP Codon | pUPD | Ampicillin | #118057 |
| | pNLuc | NLuc Luciferase | CDS + STOP Codon | pUPD3 | Chloramphenicol | #118058 |
| | pRenilla | Renilla Luciferase | CDS + STOP Codon | pUPD3 | Chloramphenicol | #118059 |
| | pGrRenilla | GrRenilla Luciferase | CDS + STOP Codon | pUPD3 | Chloramphenicol | #118060 |
| | pbGH | Bovine growth hormone terminator | 3’ UTR and poly(A) signal | pUPD3 | Chloramphenicol | #118061 |

| | | | | | | |
|---|---|---|---|---|---|---|
| Assembled Transcriptional Units and Composites | hCMV-IE1:ELuc | Constitutively expressed ELuc | Transcriptional Unit | pColE1_Alpha2 | Kanamycin | #118062 |
| | hCMV-IE1:FLuc | Constitutively expressed FLuc | Transcriptional Unit | pColE1_Alpha2 | Kanamycin | #118063 |
| | hCMV-IE1:RedF | Constitutively expressed RedF | Transcriptional Unit | pColE1_Alpha2 | Kanamycin | #118064 |
| | hCMV-IE1:NLuc | Constitutively expressed NLuc | Transcriptional Unit | pColE1_Alpha2 | Kanamycin | #118065 |
| | hCMV-IE1:Renilla | Constitutively expressed Renilla | Transcriptional Unit | pColE1_Alpha2 | Kanamycin | #118066 |
| | hCMV-IE1:GrRenilla | Constitutively expressed GrRenilla | Transcriptional Unit | pColE1_Alpha2 | Kanamycin | #118067 |
| | p(A)$^{n}$-PAUSE | Synthetic polyadenylation signal & RNA polymerase II transcriptional pause signal from the human α2 globin gene | Transcription Blocker | pColE1_Alpha1 | Kanamycin | #118068 |
| | MLRV | Multi-luciferase reporter vector | Multigenic vector | pColE1_Alpha2 | Kanamycin | #118069 |

**Supplementary Table 4. Summary of vectors used in this study.**

| Vector | Size (bp) | Molecular Weight (Da) | Transfected (ng) |
|---|---|---|---|
| **Multi-pathway luciferase reporter vector** | 13,383 | $8.28x10^6$ | 150 |
| **TB:5xNF-κβ:RedF:bGH** | 4,384 | $2.71x10^6$ | 49 |
| **TB:4xTGF-β:FLuc:bGH** | 4,394 | $2.72x10^6$ | 49 |
| **TB:5xE-box:Renilla:bGH** | 3,656 | $2.26x10^6$ | 41 |
| **TB:2xp53:NLuc:bGH** | 3,252 | $2.01x10^6$ | 36.5 |
| **TB:6xAP-1::GrRenilla:bGH** | 4,384 | $2.71x10^6$ | 41 |
| **hCMV-IE1:ELuc:bGH** | 4,789 | $2.96x10^6$ | 53.5 |

**Supplementary Table 5. DNA quantities used in experiments comparing cotransfection to solotransfection.** To compare luminescence recordings between the cotransfection and solotransfection procedures, six plasmids encoding individual luciferase transcriptional units (for cotransfection) or the multi-luciferase reporter vector (for solotransfection) were transfected in cell lines as shown in **Figure 4**. To ensure that equivalent amounts of transcriptional units for each luciferase were transfected, the number of molecules for each plasmid was calculated. For solotransfection, 150 ng of multi-luciferase reporter vector DNA was routinely transfected. Based on the molecular weight of the vector ($8.28x10^6$ Da), this corresponded to $1.09 \times 10^{10}$ molecules. This quantity was subsequently used to calculate the amount of DNA (in ng) needed to co-transfect each of the six individual plasmids.

| Pathway | Target | siRNA sequence | Reference |
|---|---|---|---|
| **p53** | TP53 | GAAAUUUGCGUGUGGAGUAdTdT | SIGMA SASI-Hs01-00056396 |
| **TGF-β** | SMAD2 | AACAAACCAGGUCUCUUGAUGdTdT | [17] |
| **NF-κβ** | NFKB1/p50 | GUCACUCUAACGUAUGCAAdTdT | [1] |
| **NF-κβ** | RELA/p65 | GAUUGAGGAGAAACGUAAAdTdT | [1] |
| **c-Myc** | MYC | GGUCAGAGUCUGGAUCACCdTdT | [18] |
| **MAPK/JNK** | JUN | AAGAACGUGACAGAUGAGCAGdTdT | [19] |
| **MAPK/JNK** | FOS | GAAUUAACCUGGGUGCUGGAdTdT | SIGMA SASI_Hs01_00184574 |
| | | CUGUCAACGCGCAGGACUUdTdT | SIGMA SASI_Hs01_00184572 |
| | | GGUUCAUUAUUGGAAUUAAdTdT | SIGMA SASI_Hs02_00184573 |

**Supplementary Table 6. Specific siRNAs used to knockdown gene expression in candidate cellular pathways.** Pathway: cellular pathway targeted. Target: mRNA transcript targeted. siRNA sequence: ribonucleotide sequence of the siRNA. Reference: publication or corporate source of sequence information. The silencing effect of each siRNA on their targets was verified by qPCR, as shown in **Supplementary Figure 15**.

| Value | # | Fold-change formula | Log2 fold-change formula | Percent change formula |
|---|---|---|---|---|
| Control value (a.u.) | 3 | $\frac{\text{Experimental value}}{\text{Control value}}$ | $\text{Log}_2\left(\frac{\text{Experimental value}}{\text{Control value}}\right)$ | $\left(\frac{\text{Experimental value}}{\text{Control value}} - 1\right) \text{x } 100\%$ |
| | 18 | 6 | 2.58 | 500% |
| | 15 | 5 | 2.32 | 400% |
| | 12 | 4 | 2 | 300% |
| | 9 | 3 | 1.58 | 200% |
| | 6 | 2 | 1 | 100% |
| | 3 | 1 | 0 | 0% |
| | 2.7 | 0.9 | -0.15 | -10% |
| | 2.4 | 0.8 | -0.32 | -20% |
| Experimental value (a.u.) | 2.1 | 0.7 | -0.51 | -30% |
| | 1.8 | 0.6 | -0.74 | -40% |
| | 1.5 | 0.5 | -1 | -50% |
| | 1.2 | 0.4 | -1.32 | -60% |
| | 0.9 | 0.3 | -1.74 | -70% |
| | 0.75 | 0.25 | -2 | -75% |
| | 0.6 | 0.2 | -2.32 | -80% |
| | 0.3 | 0.1 | -3.32 | -90% |
| | 0.15 | 0.05 | -4.32 | -95% |

**Supplementary Table 7. Ratios between experimental and control values are reported as $log_2$ fold-change.** Direct comparison between the same values calculated as fold-change, $log_2$ fold-change, and percent change.

| Cell line | ATCC number | Cancer | Media | Transfection | |
|---|---|---|---|---|---|
| | | | | Plate | Cells/well |
| **MCF7** | HTB-22 | Breast | Dulbecco's Modified Eagle Medium (DMEM) + 4 mM L-Glutamine w/o HEPES, w/o Sodium Pyruvate | 48 well | 50,000 |
| **MDA-MB-231** | HTB-26 | Breast | Dulbecco's Modified Eagle Medium (DMEM) + 4 mM L-Glutamine w/o HEPES, w/o Sodium Pyruvate | 96 well | 12,500 |
| **SK-BR-3 [SKBR3]** | HTB-30 | Breast | McCoy's 5A + 1.5 mM L-Glutamine, w/o HEPES, w/o Sodium Pyruvate | 96 well | 25,000 |
| **ZR-75-1** | CRL-1500 | Breast | RPMI 1640 + 10 mM HEPES + 1 mM Sodium Pyruvate, 4500 mg/L Glucose | 48 well | 40,000 |
| **MDA-MB-157** | HTB-24 | Breast | Dulbecco's Modified Eagle Medium/Nutrient Mixture F-12 (DMEM/F12) + 2.5 mM L-Glutamine + 15 mM HEPES | 48 well | 40,000 |
| **A549** | CCL-185 | Lung | Dulbecco's Modified Eagle Medium/Nutrient Mixture F-12 (DMEM/F12) + 2.5 mM L-Glutamine + 15 mM HEPES | 96 well | 25,000 |
| **293T/17 [HEK 293T/17]** | CRL-11268 | Non-cancerous | Dulbecco's Modified Eagle Medium (DMEM) + 4 mM L-Glutamine w/o HEPES, w/o Sodium Pyruvate | 96 well | 20,000 |

**Supplementary Table 8. Summary of cell lines with their growth and transfection conditions.** Cell culture media were supplemented with 10% US-sourced heat-inactivated fetal bovine serum (FBS) and 1% penicillin-streptomycin. All tissue culture media and supplements were obtained from Thermo Fisher Scientific. All cell lines were authenticated at the MD Anderson Characterized Cell Line Core Facility (**Supplementary Table 9** for details).

| Sample_Name | AMEL | CSF1PO | D13S317 | D16S539 | D5S818 | FGA | TH01 | TPOX | vWA |
|---|---|---|---|---|---|---|---|---|---|
| MCF7 | X | 10 | 11 | 11,12 | 11,12 | 23,24,25 | 6 | 9,12 | 14,15 |
| MCF7-Database_NCI | X | 10 | 11 | 11,12 | 11,12 | 23,25 | 6 | 9,12 | 14,15 |

| Sample_Name | AMEL | CSF1PO | D13S317 | D16S539 | D5S818 | FGA | TH01 | TPOX | vWA |
|---|---|---|---|---|---|---|---|---|---|
| MDA-MB-231 | X | 13 | 13 | 12 | 12 | 22,23 | 7,9.3 | 8,9 | 15,18 |
| MDA-MB-231-Database_NCI | X | 12,13 | 13 | 12 | 12 | 22,23 | 7,9.3 | 8,9 | 15,18 |

| Sample_Name | AMEL | CSF1PO | D13S317 | D16S539 | D5S818 | FGA | TH01 | TPOX | vWA |
|---|---|---|---|---|---|---|---|---|---|
| SK-BR-3 | X | 12 | 11 | 9 | 9,12 | 20 | 8,9 | 8,11 | 17 |
| SK-BR-3-Public Database_DSMZ | X | 12 | 11,12 | 9 | 9.,12 | 20 | 8,9 | 8,11 | 17 |

| Sample_Name | AMEL | CSF1PO | D13S317 | D16S539 | D5S818 | FGA | TH01 | TPOX | vWA |
|---|---|---|---|---|---|---|---|---|---|
| ZR-75-1 | X | 10,11 | 9 | 11 | 13 | 20,22 | 7,9.3 | 8 | 16,18 |
| ZR-75-1-Public Database_ATCC | X | 10,11 | 9 | 11 | 13 | 20,22 | 7,9.3 | 8 | 16,18 |

| Sample_Name | AMEL | CSF1PO | D13S317 | D16S539 | D5S818 | FGA | TH01 | TPOX | vWA |
|---|---|---|---|---|---|---|---|---|---|
| MDA-MB-157 | X | 10 | 11 | 11 | 12 | 22,23 | 7,8 | 9,11 | 15 |
| A549-Database_ATCC | X | 10 | 11,12 | 11 | 12 | 22 | 7,8 | 9,11 | 15 |

| Sample_Name | AMEL | CSF1PO | D13S317 | D16S539 | D5S818 | FGA | TH01 | TPOX | vWA |
|---|---|---|---|---|---|---|---|---|---|
| A549 | X,Y | 10,12 | 11 | 11,12 | 11 | 23 | 8,9.3 | 8,11 | 14 |
| A549 - Database_CLS | X,Y | 10,12 | 11 | 11,12 | 11 | 23 | 8,9.3 | 8,11 | 14 |

| Sample_Name | AMEL | CSF1PO | D13S317 | D16S539 | D5S818 | FGA | TH01 | TPOX | vWA |
|---|---|---|---|---|---|---|---|---|---|
| HEK293T/17 | X | 11,12 | 12 | 9,13 | 8,9 | 23 | 7,9.3 | 11 | 16,19 |
| HEK293T/17-Database_ATCC | X | 11,12 | 12,14 | 9,13 | 8,9 | 23 | 7,9.3 | 11 | 16,19 |

**Supplementary Table 9. Short tandem repeat (STR) profile of cell lines.** STR analysis was performed at the MD Anderson Characterized Cell Line Core Facility. This analysis verified that the STR profile of the cell lines used in this study matched the profile stored in publicly available databases.

| Pathway | Gene | Accession # | Forward primer (5'→3') | Reverse primer (5'→3') | Size (bp) | Reference |
|---|---|---|---|---|---|---|
| Housekeeping | GAPDH | NM002046 | ATGGGGAAGGTGAAGGTCG | GGGGTCATTGATGGCAACAATA | 108 | [20] |
| | CCSER2 | AK024324 | GACAGGAGCATTACCACCTCAG | CTTCTGAGCCTGGAAAAAGGGC | 143 | This work |
| | SYMPK | Y10931 | CTTCACCAAGGTTGTGCTGGAG | GCGCTTGAAGATCAGGTCTCGA | 130 | This work |
| | B2M | NM004048 | TGCTGTCTCCATGTTTGATGTATCT | TCTCTGCTCCCCACCTCTAAGT | 86 | [21] |
| | HPRT1 | M31642 | TGACACTGGCAAAACAATGCA | GGTCCTTTTCACCAGCAAGCT | 94 | [22] |
| p53 | TP53 | X02469 | GCCCAACAACACCAGCTCCT | CCTGGGCATCCTTGAGTTCC | 140 | [23] |
| | CDKN1A | S67388 | ATGGAACTTCGACTTTGTCACC | AGGCACAAGGGTACAAGACAGT | 220 | [24] |
| | BAX | NM138763 | CCCGAGAGGTCTTTTTCCGAG | CCAGCCCATGATGGTTCTGAT | 155 | [25] |
| TGF-β | SMAD2 | U59911 | ACCGAAATGCCACGGTAGAA | TGGGGCTCTGCACAAAGAT | 123 | [26] |
| | SMAD7 | AH011391 | CAGTTACCCCATCTTCATC | CATAAACTCGTGGTCATTG | 151 | [27] |
| | DAPK1 | BC143733 | CCACCACGATAGGCATGTTG | TCAAGACAGGCACGGCAAT | 68 | [28] |
| NF-κβ | RELA | M62399 | CTGCAGTTTGATGATGAAGA | TAGGCGAGTTATAGCCTCAG | 183 | [1] |
| | NFKB1 | M55643 | GTGCAGAGGAAACGTCAGAA | GTGGGAAGCTATACCCTGGA | 148 | [1] |
| | IL6 | NM000600 | ACTCACCTCTTCAGAACGAATTG | CCATCTTTGGAAGGTTCAGGTTG | 149 | [29] |
| | CCL2 | M24545 | CCCCAGTCACCTGCTGTTAT | TGGAATCCTGAACCCACTTC | 171 | [30] |
| | BCL2L1 | Z23115 | GATCCCCATGGCAGCAGTAAAGCAAG | CCCCATCCCGGAAGAGTTCATTCACT | 164 | [31] |
| c-Myc | MYC | V00568 | AATGAAAAGGCCCCCAAGGTAGTTATCC | GTCGTTTCCGCAACAAGTCCTCTTC | 112 | [32] |
| | E2F1 | NM005225 | CATCCCAGGAGGTCACTTCTG | GACAACAGCGGTTCTTGCTC | 145 | [33] |
| | TERT | NM198253 | TCACGGAGACCACGTTTCAAA | TTCAAGTGCTGTCTGATTCCAAT | 94 | [34] |
| MAPK/JNK | JUN | J04111 | CAGGTGGCACAGCTTAAACA | GTTTGCAACTGCTGCGTTAG | 80 | [35] |

| | | | | | |
|---|---|---|---|---|---|
| FOS | V01512 | AGAATCCGAAGGGAAAGGAA | CTTCTCCTTCAGCAGGTTGG | 150 | [35] |
| MMP1 | NM001145938 | AGCTAGCTCAGGATGACATTGATG | GCCGATGGGCTGGACAG | 78 | [36] |
| VEGFD | NM004469 | GTATGAACACCAGCACCTC | GGCAAGCACTTACAACCT | 121 | This work |

**Supplementary Table 10. Primers used for qPCR in this study.**

**References - Supplementary Material**
